# Greenness-Assessed RP-HPLC Method for Budesonide Quantification in Layer-by-Layer Polymeric Nanoparticles: Validation, Stability, and Controlled Release in Colonic Delivery

**DOI:** 10.1101/2025.08.11.669719

**Authors:** Giriprasath Ramanathan, Masroora Hassan, Yury Rochev

## Abstract

Robust and sustainable analytical tools are essential for the design, manufacture, and evaluation of advanced drug delivery systems. This study reports the first validated reversed-phase high-performance liquid chromatography (RP-HPLC) method for quantifying budesonide encapsulated within poly(lactic-co-glycolic acid) (PLGA)-based, layer-by-layer (LbL) coated nanoparticles intended for colonic delivery in inflammatory bowel disease (IBD). Using response surface methodology with a central composite design, optimal conditions were identified as acetonitrile:water (80:20, v/v) under isocratic mode, achieving complete separation within 5 min at a flow rate of 0.34 mL/min and detection at 244 nm. The method is buffer-free, rapid, and solvent-efficient, resulting in a favorable greenness profile, further confirmed by GAPI and NEMI assessments. Validation in line with ICH Q2(R2) demonstrated excellent linearity (R² > 0.999), precision (%RSD < 2 %), accuracy, specificity, and sensitivity (LOD: 0.04 µg/mL; LOQ: 1.2 µg/mL). Forced degradation studies under acidic, alkaline, oxidative, thermal, photolytic, and photostatic conditions showed that the LbL coating markedly enhanced drug stability, particularly against thermal and photostatic stress, while >75% degradation occurred in alkaline and oxidative environments. In vitro release profiling under simulated gastrointestinal conditions demonstrated sustained, pH-responsive release (20.6% over 48 h), consistent with colonic targeting. This validated, green, and stability-indicating method integrates controlled release assessment with comprehensive performance evaluation, providing a versatile platform for quality control of nanoparticulate drug delivery systems and supporting their progression from formulation development to clinical translation

## 1. Introduction

Inflammatory bowel disease (IBD), encompassing conditions like Crohn’s disease and ulcerative colitis, continues to pose a considerable clinical challenge. Its chronic, relapsing course, coupled with persistent difficulties in achieving sustained, targeted drug delivery, makes effective management notably complex [1]. Standard oral formulations often get absorbed too early in the upper gastrointestinal (GI) tract, which can lead to notable side effects [2], This highlights the need for colon-targeted delivery systems that allow for localized drug release and help reduce systemic exposure.[3]. To address these limitations, researchers have widely investigated nanoparticle (NPs)-based drug delivery systems for their capacity to safeguard drugs during GI transit and enable targeted release at inflamed sites. Notably, layer-by-layer (LbL) coated poly(lactic-co-glycolic acid) (PLGA) NPs have emerged as a leading approach for colonic targeting, owing to their customizable architecture, surface charge tunability, and resilience to the pH and enzymatic fluctuations encountered throughout the GI tract [4]. These systems are distinguished by their modular design, which permits surface charge change and incorporating multiple functional layers. Such features enhance both their retention and responsiveness within the colonic environment. [5,6].

Budesonide (BUd) exhibits potent topical anti-inflammatory effects as a glucocorticoid and undergoes significant first-pass metabolism, making it well-suited for localized IBD therapy. Beyond IBD, BUd has also been employed in treating various clinical conditions, including asthma [7], allergic rhinitis [8], and eosinophilic esophagitis [9], owing to its potent local anti-inflammatory action and favourable pharmacokinetic profile. It is subject to premature absorption and degradation in the upper gastrointestinal tract, which restricts its effective delivery to the colon. [10]. The therapeutic success of BUd in ulcerative colitis hinges on overcoming GI barriers through the development of advanced colon-targeted delivery systems that enable localized, controlled release at the site of inflammation [11]. Although advances have been made in formulation strategies, accurately quantifying BUd in these complex NPs systems is still quite challenging. Standard reversed-phase high-performance liquid chromatography (RP-HPLC) methods, which work fine for biological fluids or conventional dosage forms, are often unsuitable for nanoparticulate matrices. This is mainly because they rely on buffered mobile phases such as sodium dihydrogen phosphate (pH 3.2; 0.02 M) [10,12], phosphate buffer (pH 3.2–0.02 M) [13–16], ammonium formate buffer (pH 4.2, 5 Mm) in 0.1% formic acid [17], ammonium acetate (pH 3.2, 10 mM) [18,19], gradient elution [20–24], and extended run time [15], making them inefficient or unsuitable for analysing polymeric NPs systems. **(Table 1)**. Most of these methods have not been validated for use with nanoparticle matrices, particularly when subjected to forced degradation or during in vitro release studies.

**Table 1.** Reported analytical methods for BUd across different matrices, highlighting column types, mobile phases, conditions, and applications.

| Ref. | Column | Mobile Phase | Flow rate (mL/min) | Temp (°C) | Detection (nm) | Sample | Analysis | IV (µL) |
| --- | --- | --- | --- | --- | --- | --- | --- | --- |
| [10] | C18 (150×4.6 mm, 3 µm) | EtOH:ACN:Buffer (2:34:66, pH 3.2) | 1 | 50±2 | 240 | BUd | Impurity and stress degradation | 100 |
| [12] | C18 (150×4.6 mm, 5 µm) | ACN:Buffer (40:60, pH 3.5) | 2 | 25±2 | 230 | BUd as nasal sprays | Validation and Green assessment | 20 |
| [13] | C8 (150×4.6 mm) | ACN:Buffer (55:45, pH 3.2) | 1.1 | 25±2 | 244 | BUd in pectin microparticles | Stability, quantification and stress degradation | 10 |
| [14] | C18 (250×4.6 mm, 5 µm) | ACN:Buffer (55:45, pH 3.2) | 1 | 25±2 | 244 | BUd-colon targeted | Validation, release | 50 |
| [15] | C18 (150×4.6 mm, 5 µm) | EtOH:ACN:Buffer (2:30:68, pH 3.4) | 1.5 | 25±2 | 240 | Propylene glycol funtionalized BUd | Validation, Stability, and stress degradation | 10 |
| [16] | C8 (150×4.6 mm) | ACN:Buffer (32:68, pH 3.2) | 1.1 | 25±2 | 244 | BUd in rat organs | Quantification | 50 |
| [17] | C8 (150×4.6 mm, 3.5 µm) | ACN:Buffer (60:40, pH 4.2) | - | - | - | BUd in human plasma | Analytical validation | 5 |
| [18] | C18 (50×4.6 mm, 3 µm) | ACN:Buffer (35:65, pH 3.2) | 1 | 25±2 | MS/MS | BUd in human plasma | Pharmacokinetic and quantification. | 100 |
| [19] | C18 (100×3 mm, 3.5 µm) | ACN:Buffer (40:60, pH 3.1) | 1 | 25±2 | 214 | BUd | Validation and stress degradation | 10–100 |
| [20] | C18 (75×4.6 mm, 3.5 µm) | MeOH:H<sub>2</sub>O (69:31) | 0.8 | 25±2 | 254 | BUd in skin layers | Quantification | 100 |
| [21] | C18 (50×4.6 mm, 5 µm) | MeOH:H<sub>2</sub>O (70:30) | 0.8 | 25±2 | MS/MS | BUd in human plasma | Validation, stability and pharmacokinetic | 20 |
| [22] | C18 (250×4.6 mm, 5 µm) | MeOH:H<sub>2</sub>O (72:28) | 0.8 | 25±2 | 254 | BUd in rat plasma | Pharmacokinetic | 10 |
| [23] | C18 (2.1×150 mm, 5 µm) | H<sub>2</sub>O:ACN (30:70) | 1 | 30±2 | 245 | BUd in Env. water | Photo and stress degradation | NA |
| [24] | RP18 (250×4.6 mm, 5 µm) | ACN:H<sub>2</sub>O (55:45) | 1 | 35±2 | 244 | BUd microparticles in lung | Quantification and release | 5 |

From a pharmaceutical development and regulatory standpoint, method validation [25] is far more than a routine procedure. It is fundamental for ensuring the reliability, specificity, and reproducibility of analytical results throughout the product lifecycle. Without rigorous validation, the integrity of data and subsequent decisions regarding product quality would be compromised [26]. This is especially significant for NPs-based formulations, as they can experience physical or chemical alterations during processing, storage, or passage through the GI tract. Without a robust, stability-indicating analytical method [27,28], it becomes challenging to differentiate between true drug degradation and potential artifacts introduced by the analysis itself. This ambiguity can undermine the accurate assessment of formulation performance and the true stability of the product over its shelf-life [29]. The method must not only quantify total drug content but also effectively monitor in vitro drug release profiles under simulated GI conditions. In these systems, interfering substances from the release medium, polymers, or degradation products can impact baseline separation and peak resolution. Therefore, an analytical method development followed by forced degradation studies are indispensable tools in pharmaceutical development, enabling the assessment of manufacturing changes in drug quality and comparability, despite limited regulatory guidance on their standardized design and interpretation [30,31].

Conventional chromatographic methods for BUd often rely on phosphate or ammonium buffers, gradient elution, and extended run times, all of which increase solvent consumption, waste generation, and overall environmental burden. In line with the principles of green analytical chemistry, the present method was deliberately designed to be buffer-free, rapid, and solvent-efficient. Furthermore, its greenness was transparently assessed using established metrics (GAPI and NEMI), ensuring a balance between environmental sustainability and analytical functionality [32].

In this study, we designed to validate an RP-HPLC method tailored for quantifying BUd from LbL-coated PLGA NPs. The approach was kept isocratic, buffer-free, and rapid and fully validated under ICH Q2(R2) guidelines. The method was evaluated through forced degradation and applied to in vitro drug release studies to quantify BUd under stress and release from NPs efficiently for screening formulations in colon-targeted nanoparticle systems. A schematic overview of the study design, highlighting the RP-HPLC method optimization, validation, and its application to BUd NPs analysis, is presented in **Fig. 1**. Accordingly, establishing a robust, validated RP-HPLC method for BUd is not only crucial for nanoparticle formulations but also for wider analytical and clinical applications where precise drug quantification is required.

**Figure 1.**
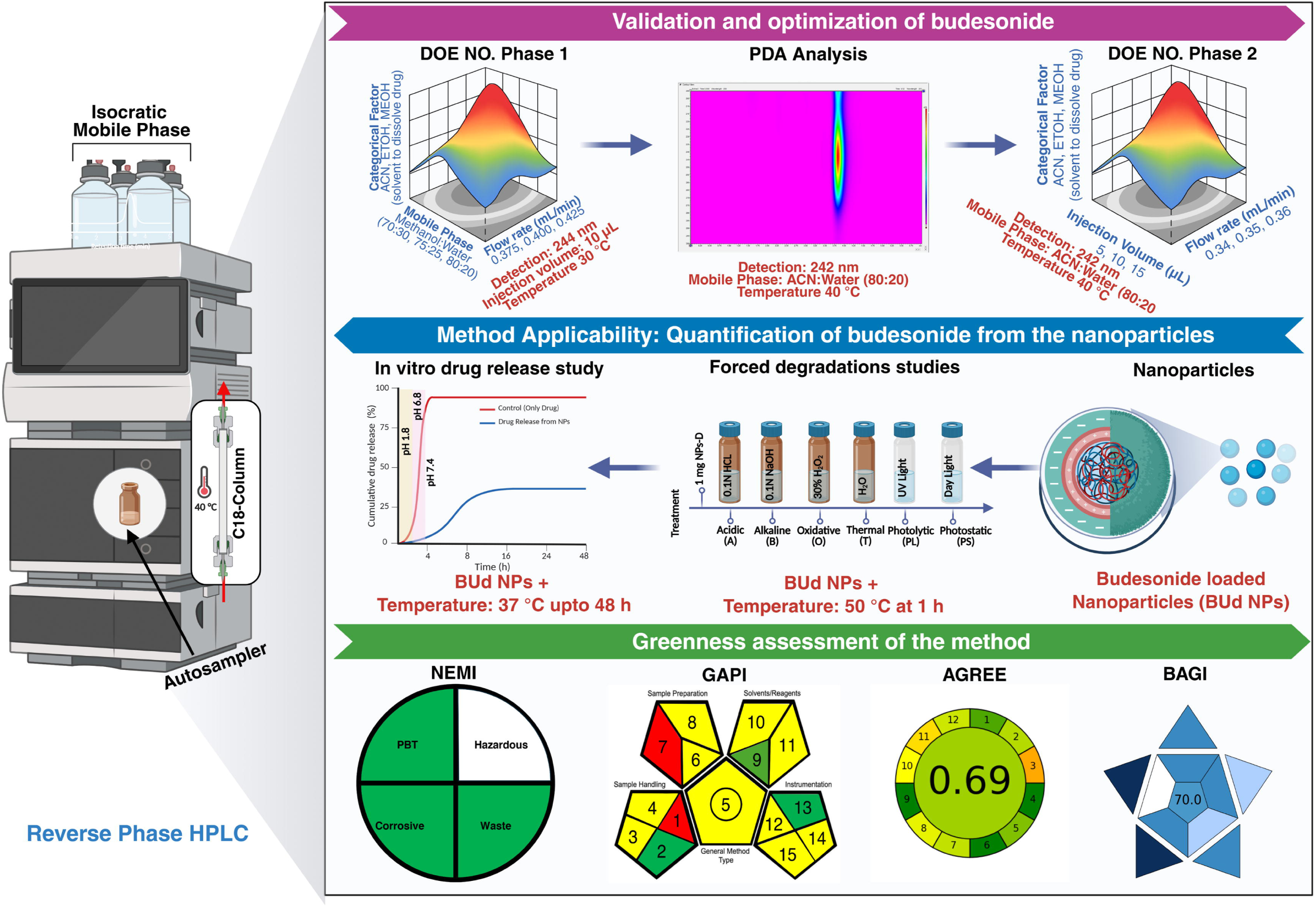
Schematic overview of the RP-HPLC method development for BUd-NPs, illustrating DOE-based optimization, method validation, and application to in vitro release and forced degradation studies.

## 2. Experimental section

### 2.1. Materials

High molecular weight hyaluronic acid (HMW-HA) (Mw 1250–1500 kDa) was purchased from Contipro (Czech Republic). It was produced biotechnologically through the fermentation of a non-haemolytic *Streptococcus* sp. strain and stored at 2–8=°C. Poly(D,L-lactide-co-glycolide) (PLGA; Resomer® RG 504 H; 50:50 lactide:glycolide ratio, Mw 38–54 kDa, amorphous) was obtained from Merck (Germany) and stored at 2–8 °C. Polyvinyl alcohol (PVA; Mw 13–23 kDa), poly-L-lysine (PLL; Mw 150–300 kDa, 0.1% w/v in water, synthetic origin), dichloromethane (DCM), sodium phosphate monobasic (MW 138.0), sodium phosphate dibasic (MW 268.0), potassium chloride (MW 74.5), hydrochloric acid, sodium hydroxide, hydrogen peroxide (H=O=), and HPLC grade solvents (acetonitrile (ACN), methanol (MeOH), ethanol (EtOH), and water (H_2_O) were also sourced from Merck. PLL was stored at room temperature. Budesonide (BUd) (MW 430.54 g/mol, ≥97 % purity) was bought from Thermo Scientific Chemicals (USA) and used as the model drug. HPLC-grade solvents were used for all analytical studies and mobile phase preparation. All other reagents and chemicals were of analytical grade and purchased from Merck unless otherwise specified.

### 2.2. Instrumentation

Chromatographic analyses were performed using a Shimadzu Prominence-i LC-2023 Plus HPLC system (Shimadzu, Japan), equipped with a quaternary pump, inline degasser, auto-sampler, column oven, sample cooler, and UV detector. Data collection, chromatogram processing, and peak integration were managed through Lab solutions software (version 5.57, Shimadzu, Japan). Experimental design, response surface modelling, and calculations involving the desirability function were performed with Design Expert® software (version 13, Stat-Ease Inc., Minneapolis, MN, USA). Particle size, polydispersity index (PDI), and zeta potential were measured by dynamic light scattering (DLS) using a Lite Sizer 500 (Anton Paar). For microstructural analysis of the particles, scanning electron microscopy (SEM) was conducted with a Hitachi S-4700 EDX.

### 2.3. Chromatographic conditions

Chromatographic separation was performed using a Kinetex® C18 column (100 × 4.6 mm, 5 µm particle size, 100 Å pore size; Phenomenex, USA) operating under isocratic conditions. Response surface methodology (RSM) with a central composite design (CCD) was used for preliminary optimization in phase I: MeOH: H_2_O mixtures (70:30, 75:25, and 80:20, v/v) served as the mobile phase. Flow rates tested included 0.375, 0.400, and 0.425 mL/min, and the column temperature was maintained at 30 °C. The injection volume was set at 10=μL throughout. UV detection occurred at a wavelength of 244=nm. Method refinement and response optimisation phase II: ACN: H_2_O (80:20, v/v) was selected as the mobile phase. The flow rates assessed were 0.34, 0.35, and 0.36 mL/min, while injection volumes of 5, 10, and 15 µL were examined. The column temperature was kept at 40 °C, and UV detection was performed at 242 nm. Across all experiments, mobile phases were filtered through polytetrafluoroethylene (PTFE) membrane filters (0.45 µm pore size, 47 mm diameter; Agilent Technologies, USA) and degassed by ultra-sonication. The auto-sampler temperature was held at 4 °C for all runs.

### 2.4. Preparation of stock solutions, calibration standards, and RSM working solutions

A primary stock solution of BUd (1000 µg/mL) was prepared by accurately weighing the drug and dissolving it in ACN. Serial dilutions were then performed using ACN and H_2_O, both filtered through 0.45 µm PTFE membranes, to obtain calibration standards at concentrations of 1 to 200 µg/mL. These standards were used in the calibration curve required for method validation. For chromatographic method optimization employing RSM-CCD, a BUd (100 µg/mL) working solution was freshly prepared and utilized in all experimental runs. To systematically assess the influence of solvent composition as a categorical factor during optimization, three organic solvents (ACN, MeOH, and EtOH) were used to dissolve BUd (100 µg/mL) prior to RSM-CCD experiments. All dilutions and preparations employed freshly prepared, degassed, and membrane-filtered (0.45 µm) to maintain consistency and ensure analytical reliability.

### 2.5. Selection and validation of key chromatographic performance parameters

The accuracy, precision, and reliability of the developed RP HPLC method were thoroughly assessed under ICH Q2(R2) guidelines [25]. Analytical performance parameters namely, peak area for quantification, retention time (Rt) for consistent analyte identification, the number of theoretical plates (N) as a reflection of column efficiency, tailing factor (T) for peak symmetry, resolution (Rs) for adequate analyte separation, and capacity factor (k’) to describe retention were systematically evaluated. Each parameter was measured against established criteria grounded in standard chromatographic benchmarks. The method’s suitability and robustness were validated comprehensively under the final chromatographic conditions, which were optimized via the second phase of RSM-CCD.

### 2.6. Experimental design and statistical analysis

#### 2.6.1. CCD for method optimisation

A design of experiments (DOE) strategy was utilized to explore the effects of key chromatographic variables on method performance, conducted in two sequential stages. These stages, phase I (screening and preliminary optimization) and phase II (method refinement and response optimization), were performed using a CCD framework within Design Expert® software (version 13, Stat-Ease Inc., Minneapolis, USA) [33,34]. Throughout both phases, Rt and PA were chosen as the primary response variables to evaluate the efficiency and sensitivity of the method.

In phase I, the composition of the mobile phase (MeOH:H_2_O) and the flow rate (mL/min) were evaluated as numerical factors, while the solvent used for drug dissolution (ACN, MeOH, EtOH) was treated as a categorical variable. In phase II, the method was further refined by adjusting both the flow rate (mL/min) and injection volume (µL), with the same categorical solvent variable.

#### 2.6.2 Model development and statistical evaluation

Each phase involved 42 experimental runs analysed using RSM. Analysis of variance (ANOVA) was employed to assess statistical significance, model adequacy, and predictive capability of the final optimised chromatographic conditions. T**able 2** summarises the independent variables and their levels used in the CCD, and the relationship between coded and actual factor values was established using the standard equation.

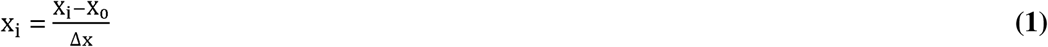

Where x_i_ is the dimensionless coded value of the i^th^ independent variable, X_i_ is the actual value of the independent variable, X_0_ is the actual value at the centre point, and Δx is the step change value of the variable. For each DOE experiment, the relationship between the independent variables and the responses (R) was modelled using the following general form:

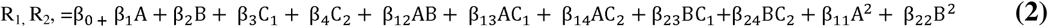

(for Phase I) and

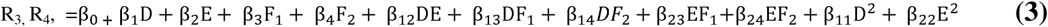

(for phase II)

Equations (2) and (3) represent the second-order polynomial models developed to predict the chromatographic responses under different experimental conditions for Phase I and Phase II, respectively. In these models, R= and R= represent Rt, while R= and R= correspond to PA, denoting the predicted responses for the two phases. The coefficients β represent the linear, interaction, and quadratic effects of the independent variables. For Phase I [Equation (2)], the independent variables A (mobile phase composition), B (flow rate), and categorical solvent variables C= and C= (ACN, MeOH, and EtOH) were used. Similarly, for Phase II [Equation (3)], the variables included D (flow rate), E (injection volume), and F= and F= (ACN, MeOH, and EtOH). The detailed levels and coding of these variables are presented in **Table 2** [34,35].

**Table 2.**
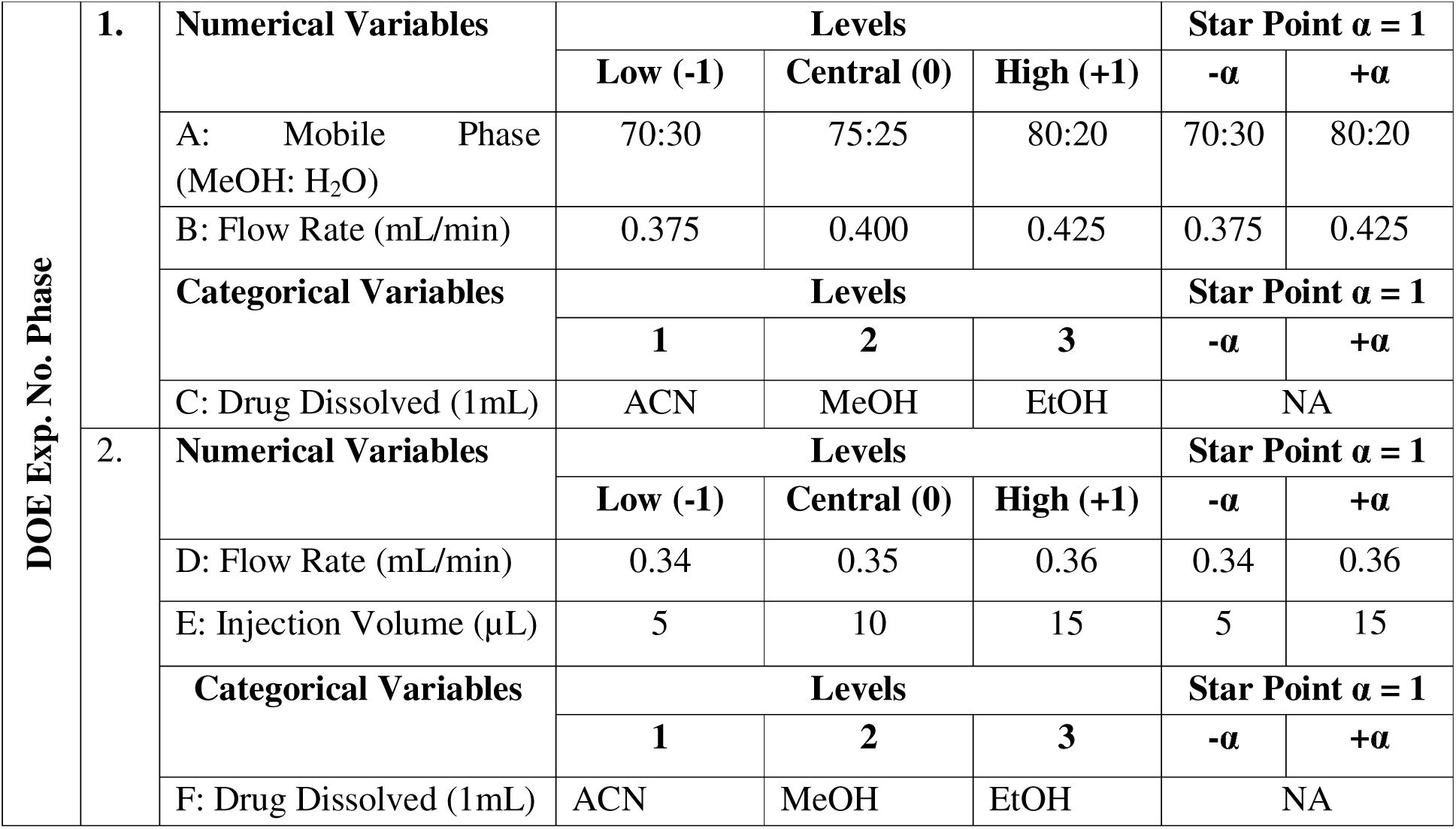
Independent variables and their corresponding levels used in RSM-CCD experiments for Phase I and Phase II method optimization.

### 2.7. Method validation parameters

#### 2.7.1. Specificity

Chromatographic conditions were optimized for specificity and selectivity to ensure unambiguous analyte quantification in the presence of excipients, degradation products, and matrix interferences. Two separate experimental strategies were employed: matrix interference and forced degradation studies.

##### 2.7.1.1. Matrix interference

Specificity was assessed by injecting a series of samples, mobile phase (blank), placebo, NPs without BUd (Bare NPs), standard BUd solution (BUd), BUd –loaded NPs (BUd NPs), and BUd NPs subjected to forced degradation. Each sample was analysed under optimized chromatographic conditions to verify the absence of interfering peaks at the BUd retention time. Peak purity was evaluated using photodiode array (PDA) detection, and contour plots (200–300 nm) were generated to confirm spectral homogeneity and rule out co-elution of degradation products. This ensured the method could selectively quantify BUd in complex nanoparticle matrices, even under stressed conditions [34,36].

##### 2.7.1.2. Forced degradation study

To assess the stability-indicating properties of the developed RP-HPLC method, forced degradation studies were conducted in line with the ICH Q1A(R2) guidelines. These experiments examined the chemical stability of both free BUd and BUd NPs under a range of stress conditions to identify possible degradation products and validate the specificity of the method. Stress conditions applied included hydrolytic (acidic and basic), oxidative, thermal, photolytic, and photostatic degradation. For each condition, the following sample groups were prepared as untreated BUd NPs as NPs control (NC), BUd NPs at 50 °C subjected to the respective stress (N50), free BUd at 50 °C subjected to the respective stress (D50), free BUd at 37 °C subjected to the respective stress (D37), BUd NPs subjected to photolytic and photostatic (NUV and NDL), and BUd subjected to photolytic and photostatic (DUV and DDL) as illustrated in **Fig. 7(a)**. Hydrolytic degradation was induced by treating samples with 0.1 N and 1 N HCl or NaOH at 50 =°C for 1 h. Oxidative degradation was performed using 30 % H=O= at 50= °C for 1 h. For thermal degradation, samples underwent dry heating at 50= °C for 1 h without additional reagents. BUd and Bud NPs were suspended in HPLC grade water in transparent glass vials were exposed to UV-C light (254=nm) at an intensity (13.46=W·h/m²) for 1 h and visible light from an ET17 tube lamp (15.8 million lux·h) for 1 h for photolytic and photostatic degradation, respectively. Control samples, in both cases, were protected from light in order to clearly isolate and assess the effects of illumination. Following stress treatments, all samples were neutralized when necessary, freeze-dried, redissolved in a defined volume of ACN, filtered through a PTFE membrane (0.22 µm), and analysed using the validated RP-HPLC method. This systematic approach to sample preparation enabled precise quantification of BUd and reliable detection of degradation products across all tested conditions [34,35].

#### 2.7.2. Linearity

Linearity was assessed by analysing standard BUd solutions across a concentration range of 1–200 µg/mL. A calibration curve was generated by plotting peak area against concentration, followed by linear regression analysis to obtain the correlation coefficient (r) and coefficient of determination (r²). These results confirmed that the method yields responses directly proportional to the analyte concentration [36].

#### 2.7.3.. Accuracy

Method accuracy was evaluated through recovery studies conducted at three concentration levels (100, 93.3, and 86.6 %) of the target concentration, which corresponded to theoretical values of 187.5, 175.0, and 162.5 µg/mL, respectively. These BUd concentrations were prepared volumetrically by mixing two stock solutions, namely vial A (200 µg/mL) and vial B (150 µg/mL), in ratios of 750:250, 500:500, and 250:750 µL. Each concentration level was analysed in triplicate. Accuracy was determined by calculating the mean recovery, standard deviation (SD), and percent relative standard deviation (% RSD). The acceptance criteria were set at 98 to 102 % recovery and ≤ 2 % RSD [14,37].

#### 2.7.4. Precision

The precision of the developed RP-HPLC method was assessed by evaluating repeatability (intra-day) and intermediate precision (inter-day). Repeatability was determined by analysing six replicates of a BUd (200 µg/mL) standard solution under identical conditions within the same day. Intermediate precision was evaluated by repeating this procedure on the subsequent day, using the same instrumentation and methodological parameters. Precision was reported as the % RSD (≤ 2 %) considered acceptable [34,35].

#### 2.7.5. Sensitivity

The limit of detection (LOD) and limit of quantification (LOQ) of the developed method were established using the signal-to-noise (S/N) approach, with approximate ratios of 3:1 and 10:1, respectively., A statistical assessment was conducted by employing the standard deviation of the response (σ) and the slope of the calibration curve (S), utilizing the following equations:

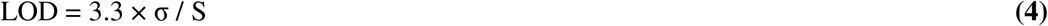

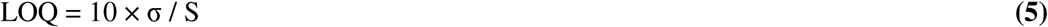

LOD indicates the lowest concentration of BUd that can be reliably detected, and LOQ corresponds to the minimum concentration that can be quantified with acceptable precision and accuracy [14,35].

#### 2.7.6. Robustness

Robustness was assessed by making small, intentional adjustments to key chromatographic parameters, in order to determine how well the method withstood typical operational changes. Parameters examined were flow rate (± 0.01 mL/min), injection volume (± 5 µL), column temperature (± 10 °C), and detection wavelength (± 2 nm). Each modified condition was analysed twice, and the effects on retention time, peak area, and overall system suitability were evaluated. Consistency in chromatographic performance across these variations was used as evidence of the method’s robustness [34].

#### 2.7.7. System Suitability

System suitability was evaluated to confirm reliable performance of the RP-HPLC system before analysing samples. Six replicate injections of a standard BUd solution (100 µg/mL) were performed under the optimized chromatographic conditions. Key chromatographic parameters as Rt, PA, number of theoretical plates (N), tailing factor (T), resolution (Rs), and capacity factor (k’) were carefully recorded. The % RSD for both peak area and retention time was calculated to assess repeatability and instrument stability [38].

### 2.8. Application of the validated RP-HPLC method to nanoparticle formulation

#### 2.8.1. Synthesis of budesonide-loaded nanoparticles (BUd NPs)

The HA-functionalized, BUd NPs were prepared using a previously established co-solvent and LbL assembly method with minor modifications to incorporate the drug was schematically illustrated in **Fig. 2(a)** [39]. Briefly, PLGA (2 % w/v) and BUd (5 mg, pre-dissolved in EtOH)) were dissolved in a 3:1 (v/v) mixture of DCM and EtOH. This organic phase was added drop-wise into 1 % (w/v) aqueous PVA under probe sonication (8=/2=s:on/off, 120=s total) to form an oil-in-water emulsion, followed by overnight stirring at room temperature for solvent evaporation and nanoparticle formation. The resulting BUd –loaded PLGA NPs (BUd-PLGA NPs) were collected by centrifugation (10,350 rpm, 15 min), washed, and subjected to sequential surface modification via electrostatic adsorption. For the first coating step, BUd-PLGA NPs were incubated in PLL (0.1 % (w/v) solution for 12 h, enabling cationic PLL chains to adsorb onto the negatively charged PLGA surface, forming BUd-PLGA-PLL NPs. These were subsequently incubated in HMW-HA (0.2 % (w/v)) solution for 4 h, allowing electrostatic attraction between the anionic HA and the cationic layer (PLL) to yield the final BUd-PLGA-PLL-HA NPs (BUd NPs). The completed NPs were lyophilized and stored under desiccated conditions until further use.

**Figure 2.**
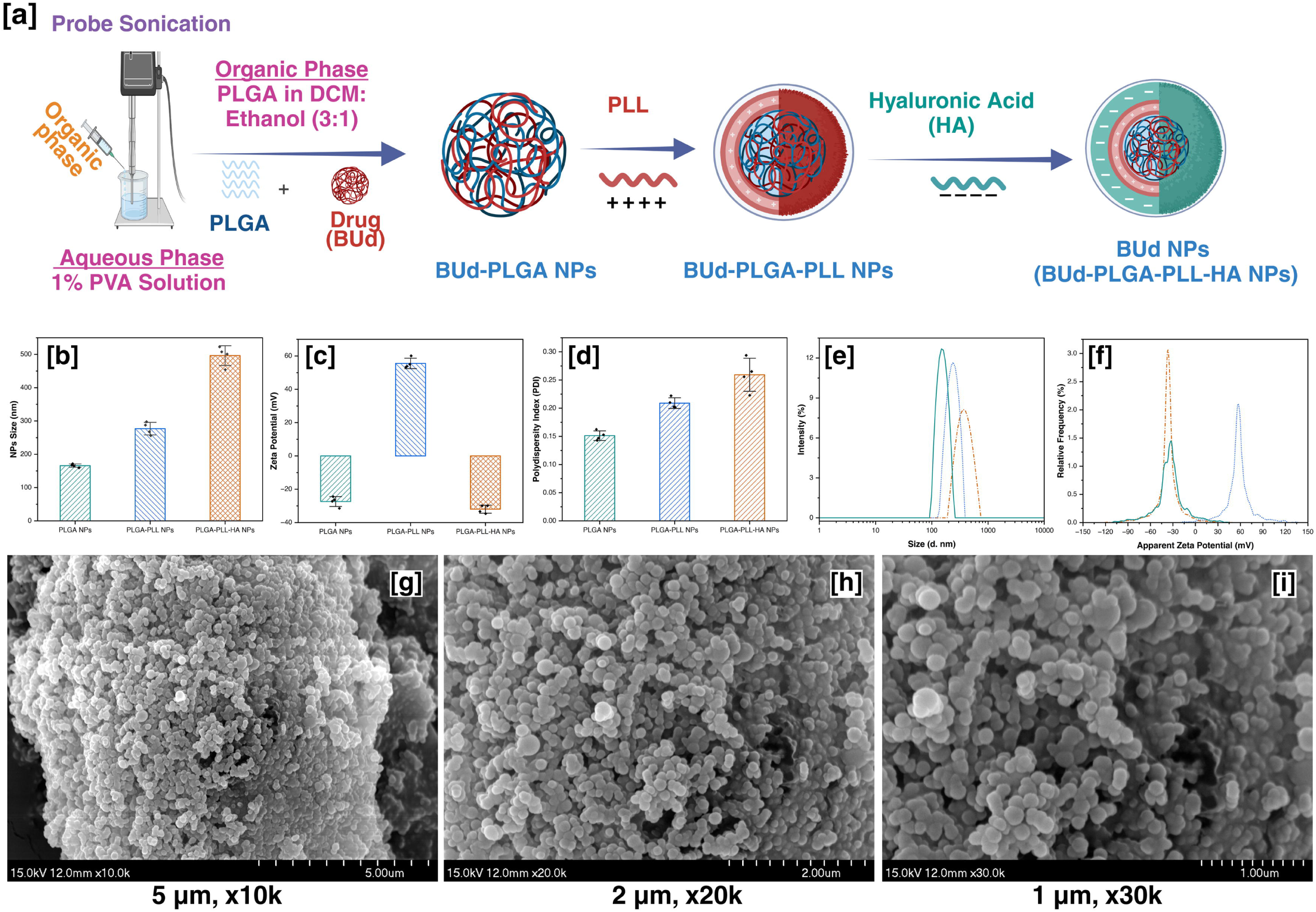
(a) Schematic representation of the sequential electrostatic coating of BUd-PLGA NPs with PLL and HA to obtain BUd NPs. Physicochemical characterization including (b) hydrodynamic diameter, (c) zeta potential, (d) PDI, (e) intensity, and (f) frequency (%) distributions by DLS (n = 3). (g–i) SEM images at 5=µm (×10k), 2=µm (×20k), and 1=µm (×30k).

#### 2.8.2. Determination of drug loading in nanoparticles

To determine drug loading, approximately 1 mg of lyophilized BUd NPs was dissolved in 1 mL of ACN. The sample was vortexed thoroughly to ensure complete disruption of the NPs and release of the encapsulated drug into the solvent. The resulting solution was then filtered using a 0.22 µm syringe filter. The clear filtrate was subsequently analysed using the validated RP-HPLC method described previously. BUd content in the nanoparticle sample was quantified by referencing a standard calibration curve. Theoretically, 1 mg of BUd NPs is expected to contain 200 µg of BUd.

#### 2.8.3. In vitro drug release studies from BUd NPs

The in vitro release profile of BUd from BUd NPs was evaluated using a dialysis-based approach that replicates the sequential pH changes of the GI. For these experiments, Slide-A-Lyzer® MINI Dialysis Devices (10 kDa MWCO, 50 mL tube) were employed, as outlined in **Fig. 8(a)**. Precisely 1 mg of lyophilized NPs was weighed, dispersed in 1 mL of HPLC grade water, and loaded into the sample chamber of each dialysis unit. The devices were immersed in 45 mL of pre-warmed release medium maintained at 37=±=0.5 °C, with constant agitation at 200 rpm. To mimic GI transit, the release study was performed under a stepwise pH regimen. Initially, the dialysis units were incubated in simulated gastric fluid (SGF, pH 1.2) for 1 h. This was followed by transfer to fresh, pre-warmed simulated intestinal fluid (SIF, pH 6.8) for 3 h. Subsequently, the medium was replaced with simulated colonic fluid (SCF, pH 7.4), and the release profile was monitored for up to 72 h. At predetermined time intervals (1, 2, 4, 8, 24, and 48 h), 1 mL samples were withdrawn from the external release medium, immediately frozen, and lyophilized. The dried samples were then reconstituted in a known volume of ACN to solubilize the released drug. Quantitative determination of BUd was achieved using a validated RP-HPLC method. To maintain sink conditions, the sampled volume was replaced with a fresh, pre-warmed buffer of the corresponding pH at each time point [4].

### 2.9. Greenness assessment tools and evaluation approach

The greenness of the developed RP-HPLC method was evaluated using three complementary tools: the National Environmental Methods Index (NEMI), the Green Analytical Procedure Index (GAPI), the Analytical GREEnness metric (AGREE) and the Blue Analytical Greenness Index (BAGI). NEMI provides a four-quadrant pictogram reflecting compliance with key criteria, namely the absence of persistent, bioaccumulative, and toxic (PBT) chemicals, non-corrosive mobile phase conditions (pH 2–12), absence of hazardous waste, and total waste volume per run (<10 mL, 10–100 mL, or >100 mL) [32]. GAPI enables a broader life-cycle evaluation of 15 parameters covering sampling, preservation, transport, storage, sample preparation, extraction type and scale, solvent selection, additional treatments, reagent amount and hazard, energy consumption, occupational risk, waste generation, and waste treatment. Each is classified on a three-color scale (green, yellow, red), and the cumulative outcome is presented as a pictogram [32]. AGREE assesses adherence to the 12 principles of green analytical chemistry using open-access software (https://mostwiedzy.pl/AGREE.). Each principle is scored from 0 to 1, and the overall average is visualized in a circular “clock” diagram, where each segment corresponds to one principle [40]. The analytical sustainability was benchmarked using the BAGI (https://bagi-index.anvil.app/), which assigns scores (10–2.5) across 10 attributes, including type of analysis, number of analytes, instrumentation, sample preparation, throughput, reagent availability, preconcentration, automation, and sample amount. Scores were summed to provide an overall value (0–100), displayed in a star-shaped radar plot [41].

### 2.10. Statistical analysis

All experiments were performed in triplicate unless otherwise stated, and results are expressed as mean ± standard deviation (SD). Calibration, linear regression, and sensitivity parameters were analysed using (OriginPro 2024, OriginLab, USA). Validation data, including accuracy, precision, robustness, and system suitability, were evaluated according to ICH guidelines. In vitro release and forced degradation studies were statistically compared using one-way ANOVA followed by Tukey’s post hoc test, with significance set at *p* < 0.05. Graphical representations include error bars for standard deviations, and all annotated comparisons reflect statistically significant differences.

## 3. Results and discussion

### 3.1. Physicochemical properties of the BUd NPs

The drug (BUd) loaded NPs displayed progressive increases in particle size and stepwise surface charge reversal, confirming successful LbL assembly. The final formulation showed a mean hydrodynamic diameter of (496 ± 30 nm) with a PDI below 0.25 ± 0.02, indicating uniform size distribution. Zeta potential shifted from (–27.4 ± 3.0 mV) (uncoated) to (55.5 ± 3.1 mV) (PLL coated), and back to (−32.0 ± 2.4 mV) after HA deposition **(Fig. 2(b-f))**. The final size remained within the submicron range with a low PDI, indicating colloidal stability of the formulation with distinct surface charge transitions, further support sequential polymer deposition and colloidal stability [4,5,39].

### 3.2. Morphological analysis of the BUd NPs

SEM analysis confirmed that BUd NPs exhibited a spherical and uniform morphology across different magnifications, with no signs of aggregation **(Fig. 2(g-i))**. The SEM morphology revealed that the BUd NPs were ranged from around 500 nm with monodisperse colloidal nature and observed particle sizes were consistent with DLS measurements [39,42].

### 3.3. Optimization of the method using RSM-CCD

#### 3.3.1. Phase I: Screening and preliminary optimization

In the initial phase, RSM-CCD was employed to screen key chromatographic factors influencing BUd quantification, specifically mobile phase composition, flow rate (numerical), and drug dissolution solvent (categorical). While MeOH-based systems provided acceptable retention and peak shape, they consistently caused higher back-pressure at elevated flow rates because of methanol’s viscosity and hydrogen bonding with water [43]. Additionally, dissolving BUd in MeOH or EtOH led to variable peak areas and poor reproducibility. In contrast, acetonitrile markedly improved BUd solubility and signal response, attributed to its lower viscosity, weaker hydrogen bonding, and stronger elution strength [44]. These experimental findings (R_1_, R_2_, and R_2a_), summarized in **Table S1**, supported the transition to an acetonitrile-based system for Phase II optimization. Second-order polynomial models (Eq. 6 and 7) were generated using Design Expert software to predict retention time and peak area in Phase I.

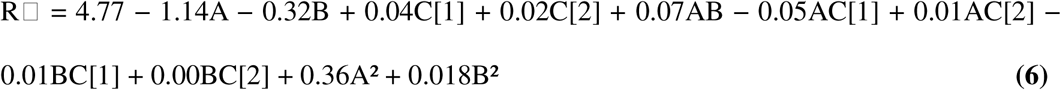

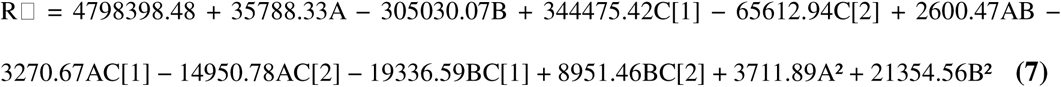

Where, A denotes mobile phase ratio, B is flow rate, and C[1] and C[2] represent solvent categories. ANOVA analysis **(Table S3)** confirmed the model’s significance with high F-values (3919.94 for Rt and 1392.43 for PA) and p-values < 0.0001, indicating robust predictability [33,34,45].

#### 3.3.2. Phase II: Method refinement and response optimization

Building on Phase I findings, Phase II focused on optimizing flow rate and injection volume under a fixed ACN:H=O (80:20, v/v) mobile phase. The aim was to improve reproducibility, signal intensity, and system stability while minimizing back-pressure. Lower flow rates reduced system strain, and a 10=µL injection volume yielded strong, undistorted signals. These trends aligned with the performance characteristics of the core-shell Kinetex C18 column (100=mm × 4.6=mm), known for high-efficiency separations at reduced pressure [44]. Using Design Expert software, second-order polynomial models (Eq. 8 and 9) were developed to predict retention time and peak area in Phase II.

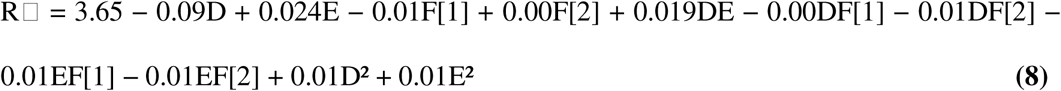

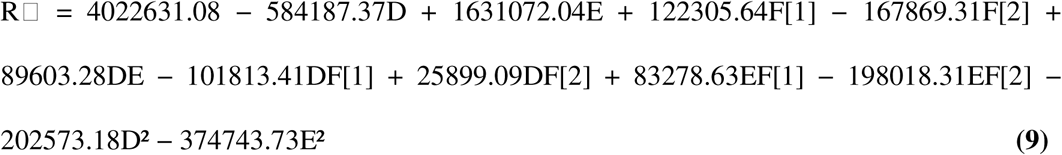

Where, D represents flow rate, E denotes injection volume, and F[1] and F[2] indicate solvent categories. ANOVA results **(Tables S4)** showed significant model fits, with F-values of 21.17 for retention time and 10.62 for peak area (p=<=0.0001). These experimental findings (R_3_, R_4_, and R_4a_), were summarized in **Table S2**.

#### 3.3.3. Influence of column temperature and PDA-based peak purity assessment

During method refinement, the effect of column temperature on detection wavelength and spectral behaviour was evaluated. At 30 °C, BUd showed maximum absorbance at 244 nm, while increasing the temperature to 40 °C caused a hypsochromic shift to 242 nm, likely due to temperature-induced changes in the chromophore environment, consistent with solvatochromic effects [44]. PDA-based peak purity analysis at both temperatures confirmed spectral homogeneity **(Fig. 5d)**, ruling out degradation or co-elution. These results highlight the importance of temperature and wavelength optimization in RP-HPLC, especially for thermosensitive or lipophilic analytes.

#### 3.3.4. Summary of optimized parameters

The method development and optimization strategy led to a finalized set of RP-HPLC conditions suitable for the quantification of BUd. The optimized method employed a mobile phase of ACN:H_2_O (80:20, v/v), a flow rate of 0.34 mL/min, and an injection volume of 10 µL. The optimal detection wavelength was determined to be 242 nm at a column temperature of 40 °C, and ACN was used as the solvent for dissolution of drug, because of its superior solubilization properties and compatibility with the mobile phase. A Kinetex C18 column (100 mm × 4.6 mm) was selected for its high efficiency and ability to operate at low back-pressure.

#### 3.3.5. Effect of variables in Phase I and Phase II

The relationship between the independent variables to understand factor interactions to get responses (R_1_, R_2_, R_3_, and R_4_) across both phases of method development using RSM-CCD was effectively generated from the empirical models (Equations 6–9). The responses (Rt and PA), for both Phase I and II are presented as two-dimensional contour plots, three-dimensional surface plots, and actual vs. predicted plots, using ACN as the categorical solvent was effectively depicted in **Fig. 3(a–f)** and **Fig. 4(a–f)**, respectively. Corresponding results for MeOH and EtOH as categorical solvents are provided in supplementary **Fig. S1 and S2**.

**Figure 3.**
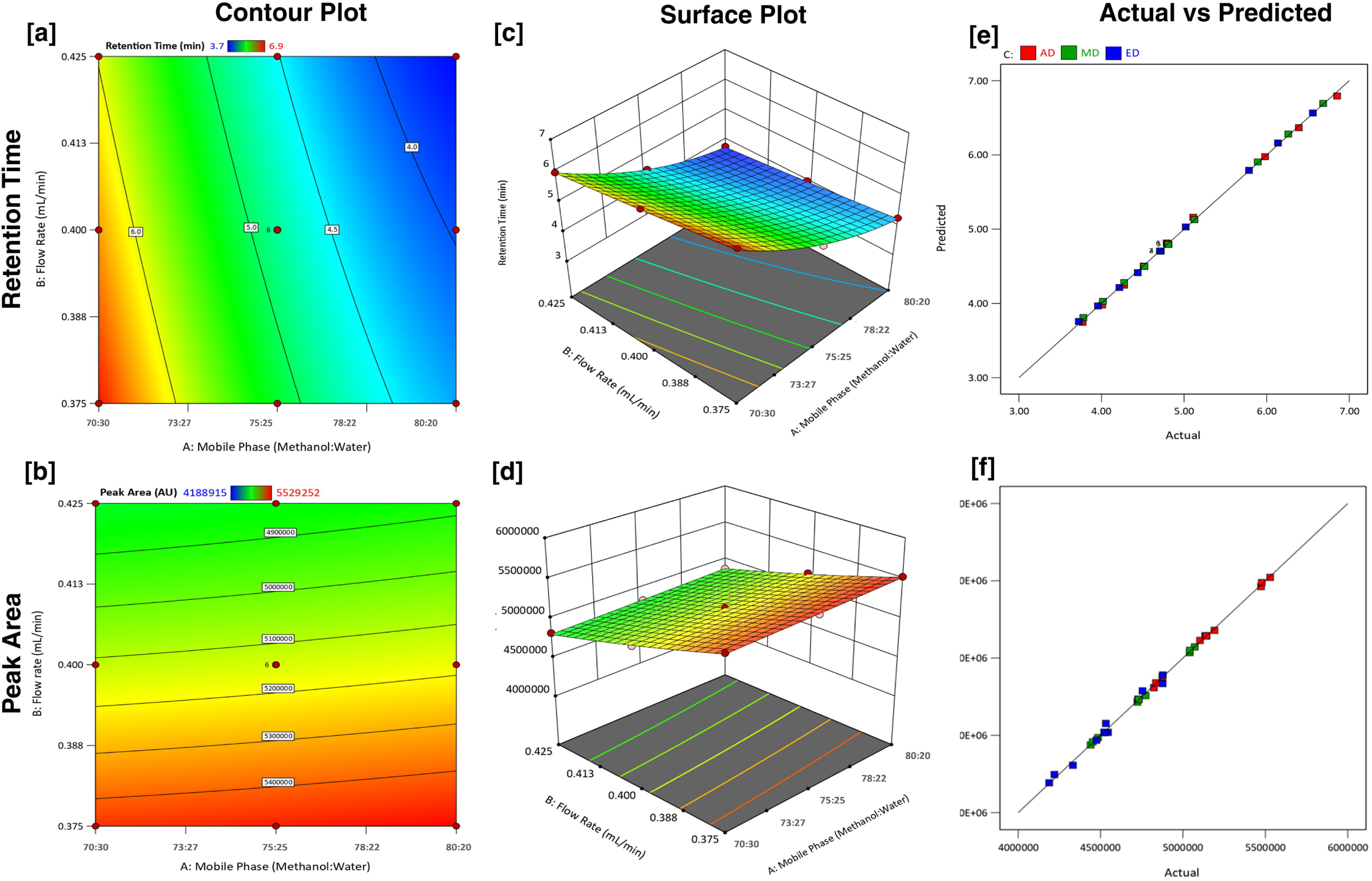
Visualization of variable interactions for phase I optimization of the RP-HPLC method with ACN as the categorical solvent. (a, b) Contour plots; (c, d) 3D surface plots; and (e, f) actual vs. predicted plots showing the combined effects of MeOH:H_2_O composition (70:30, 75:25, 80:20, v/v) and flow rate (0.375, 0.400, and 0.425=mL/min) on retention time and peak area (n = 3).

**Figure 4.**
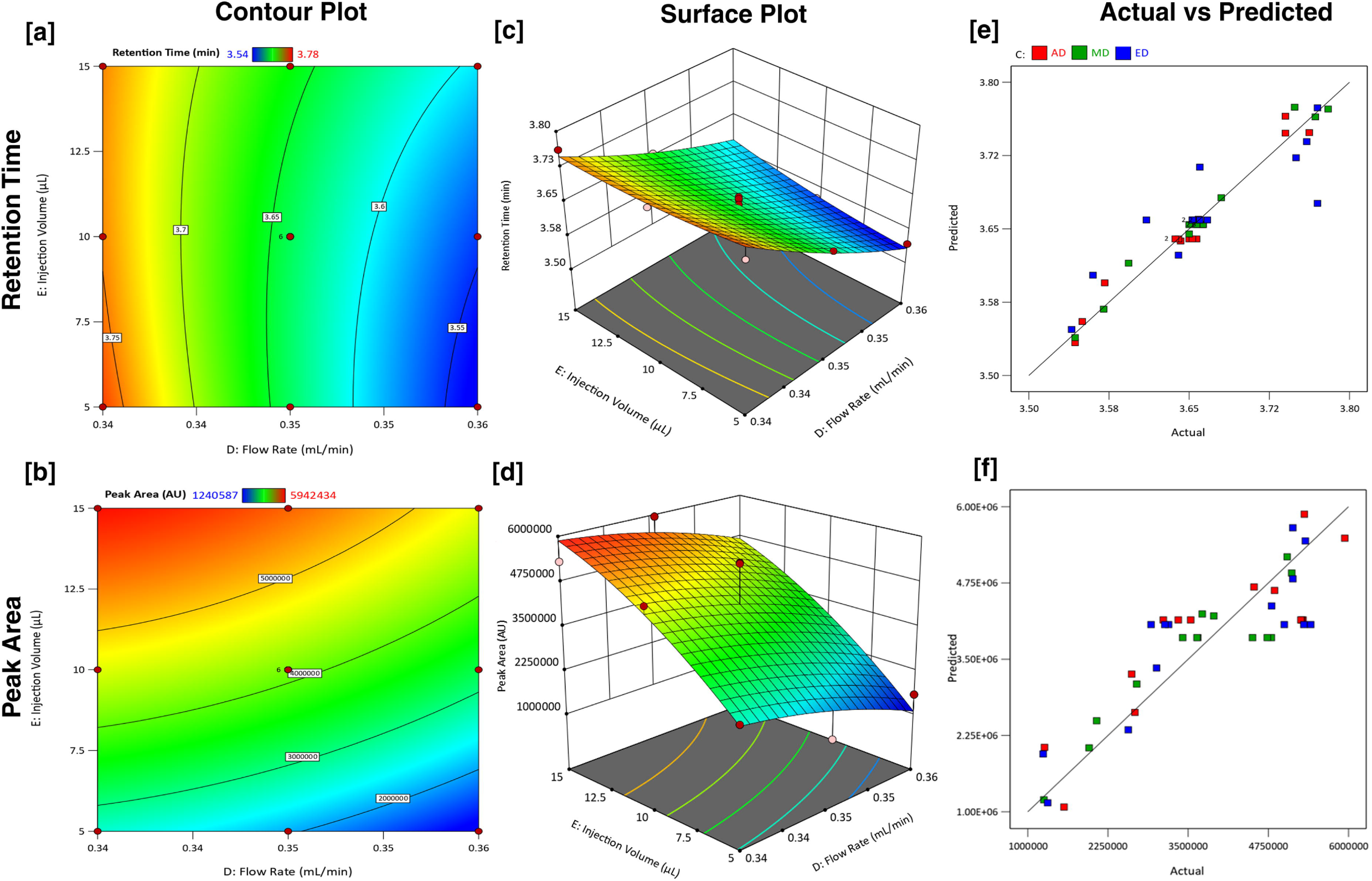
Visualization of variable interactions for phase II optimization of the RP-HPLC method with ACN as the categorical solvent. (a, b) Contour plots; (c, d) 3D surface plots; and (e, f) actual vs. predicted plots showing the combined effects of flow rate (0.34, 0.35, and 0.36=mL/min) and injection volume (5, 10, and 15=µL) on retention time and peak area (n = 3).

**Figure 5.**
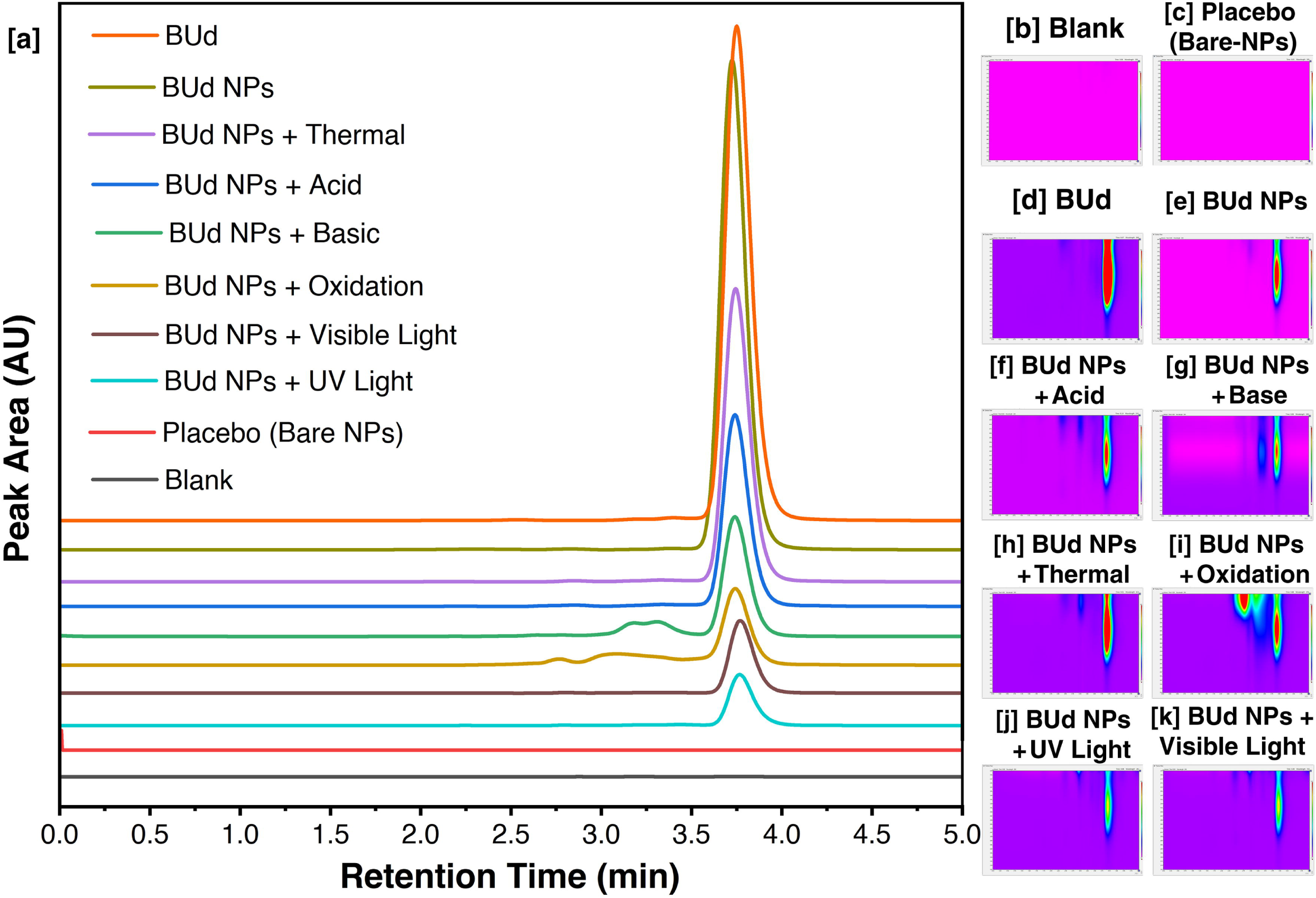
(a) Representative RP-HPLC chromatograms showing method specificity for blank, placebo, BUd, BUd NPs, and BUd NPs under stress. PDA contour plots (200– 320=nm) for (b) blank, (c) Bare NPs, (d) BUd, (e) BUd NPs, and stressed BUd NPs under (f) acidic, (g) basic, (h) thermal, (i) oxidative, (j) UV, and (k) visible light conditions.

In Phase I, both flow rate and mobile phase composition significantly affected Rt) and PA. Rt decreased with increasing flow rate and acetonitrile content, while PA peaked at intermediate flow rates with higher acetonitrile ratios. Acetonitrile showed superior solvent compatibility and system stability, supporting its selection for Phase II. The statistical robustness of the models was strong, with R²/adjusted R² values of 0.993/0.991 for Rt (Eq. 6) and 0.998/0.997 for PA (Eq. 7). ANOVA confirmed model significance with a non-significant lack-of-fit, validating the models for prediction and optimization. [33].

In Phase II, flow rate and injection volume were evaluated using an ACN:H=O (80:20, v/v) system. Injection volumes above 10 µL caused minor peak broadening, while reduced flow rates slightly improved resolution without extending runtime. Although the effects on Rt and PA were less pronounced than in Phase I, the design identified an optimal balance between signal intensity and peak shape. The model for Rt (Eq. 8) showed moderate fit (R² = 0.889; adj R² = 0.847), while the PA model (Eq. 9) yielded R² = 0.801 and adj R² = 0.725. ANOVA confirmed statistical significance, with a non-significant lack-of-fit for Rt. The slightly higher lack-of-fit in the PA model may reflect injection-volume-related variability, as previously reported in low-volume LC systems. [34]. Collectively, the model-driven findings highlighted the critical influence of mobile phase strength, flow rate, and injection volume on chromatographic performance. Acetonitrile emerged as the preferred solvent, offering superior dissolution, consistent elution, and low back-pressure. The validated models provide confidence in the optimized parameters, supporting their use in method validation and application to BUd NPs.

### 3.5. Validation of RP-HPLC method

#### 3.5.1. Specificity

The developed RP-HPLC method exhibited excellent specificity, with no interfering peaks at the BUd retention time (3.75 ± 0.01=min) in blank and placebo chromatograms **(Fig. 5a).** BUd consistently eluted as a sharp, symmetrical peak with stable Rt across all conditions. Under basic and oxidative stress, minor additional peaks indicated degradation products. PDA-based peak purity analysis confirmed spectral integrity, with purity angles below threshold and contour plots (200 – 320=nm) showing no spectral overlap, except in stressed samples, where distinct degradant peaks were observed **(Fig. 5(b) to (k)** [36]

#### 3.5.2. Linearity

The method demonstrated excellent linearity over a broad concentration range (1–200 µg/mL) **(Fig. 6(a) and (b))**, covering the expected levels for in vitro release and degradation studies. The calibration curve followed the equation *y*===20291.68 + 52847.85*x*, with a correlation coefficient (r) of 0.999 and r² of 0.998. All calibration points exhibited % RSD (< 2 %) **(Table S4)**, confirming accurate and reliable quantification across both low and high concentration ranges. [14].

**Figure 6.**
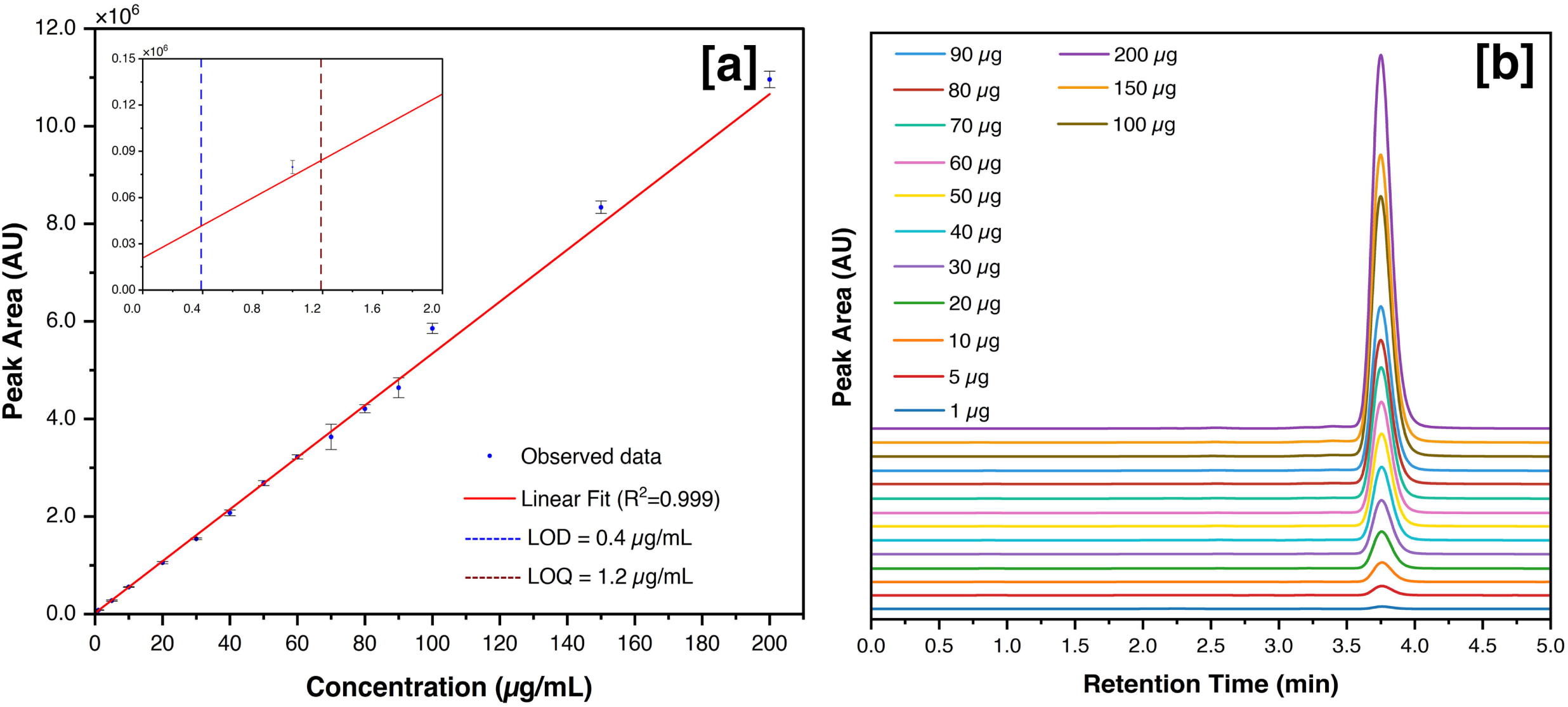
Calibration and sensitivity assessment for BUd (1–200=µg/mL). (a) Calibration curve with linear regression and inset showing LOD and LOQ values. (b) Overlaid RP-HPLC chromatograms demonstrating method sensitivity and linearity (n = 6).

**Figure 7.**
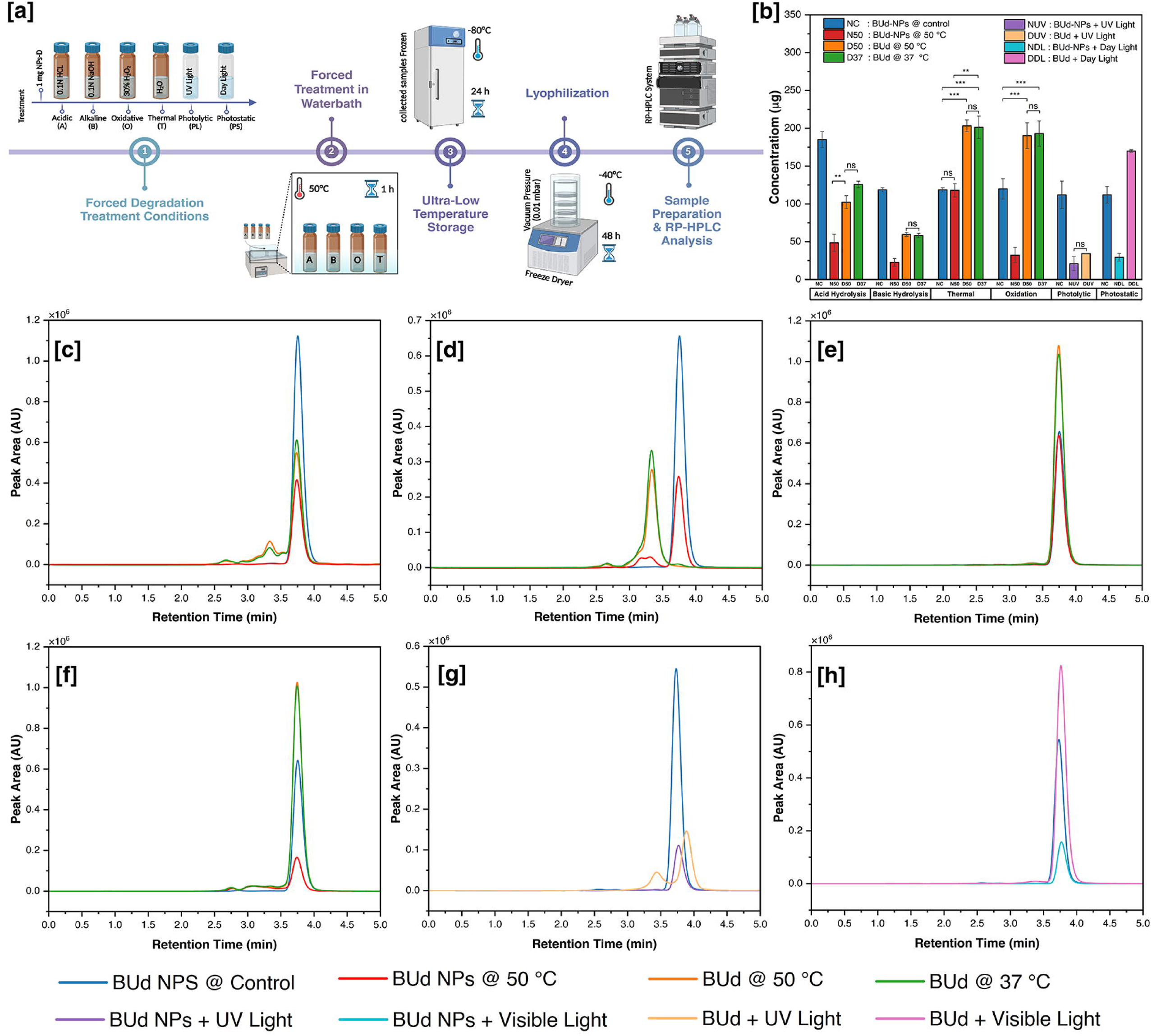
(a) Schematic of forced degradation protocol for BUd and BUd NPs under hydrolytic (acidic, basic), oxidative, thermal, photolytic (UV), and photostatic (visible light) stress. (b) Quantitative BUd content analysis in BUd NPs under stress (n = 3). ANOVA with Tukey’s post hoc test: *p=<=0.05, **p=<=0.01, ***p=<=0.001, ****p=<=0.0001. Only p ≥ 0.0001 comparisons are annotated, all others are statistically significant at p < 0.0001. (c– h) RP-HPLC chromatograms showing degradation profiles under (c) acidic, (d) basic, (e) thermal, (f) oxidative, (g) UV, and (h) visible light conditions.

**Figure 8.**
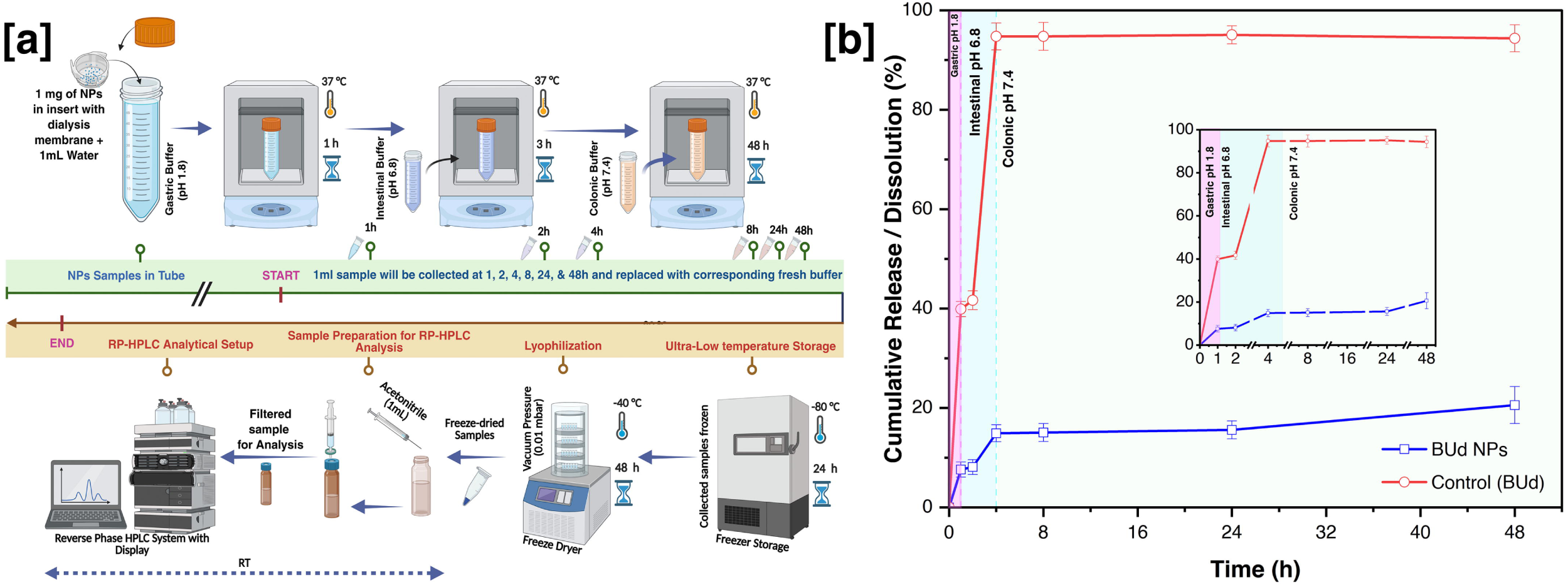
(a) Schematic of the in vitro release protocol simulating gastrointestinal pH transitions using dialysis devices (pH 1.8, 6.8, and 7.4 at 37 °C). (b) Release profile of BUd from BUd-NPs over 48 h with inset showing segmented x-axis for early-phase visualization. Data shown as mean ± SD (n = 3).

#### 3.5.3. Accuracy

Accuracy was evaluated through recovery studies at three spiking levels, yielding mean recoveries between 98.1 to 99.7 % with % RSD values from 0.8 % to 1.4 %, confirming the method’s reliability in the presence of excipients **(Table 3(a))**. These results also demonstrate that the method accurately quantifies BUd without loss or interference from sample handling or dilution steps, ensuring recovery of the pure drug even in matrix-containing samples [14,37].

**Table 3.** Summary of method validation parameters for BUd by RP-HPLC. (A) Accuracy (n===3). (B) Robustness evaluation (n===3). (C) System suitability parameters (n===6).

| (a) Accuracy |  |  |  |  |  |
| --- | --- | --- | --- | --- | --- |
| Spiking Ratio (A:B, $\mu\text{L}$) | Theoretical Value ($\mu\text{g/mL}$) | | Experimental Mean ($\mu\text{g/mL}$) | Recovery (%) | % RSD |
| 750:250 | 187.5 | | 186.9 ± 1.5 | 99.7% | 0.8% |
| 500:500 | 175.0 | | 172.5 ± 1.3 | 98.6% | 0.8% |
| 250:750 | 162.5 | | 159.4 ± 2.2 | 98.1% | 1.4% |
| (b) Robustness |  |  |  |  |  |
| Parameter | Level | Retention Time (min) |  | Peak Area |  |
| | | Mean ± SD | RSD (%) | Mean ± SD | RSD (%) |
| Flow rate ( mL/min) | 0.34 | 3.7 ± 0.0 | 0 | 4847382 ± 22402 | 0.5 |
| | 0.35 | 3.65 ± 0.01 | 0.3 | 4303177 ± 71076 | 1.7 |
| Injection Volume ($\mu\text{L}$) | 10 | 3.7 ± 0.0 | 0.0 | 4847382 ± 22402 | 0.5 |
| | 15 | 3.76 ± 0.02 | 0.5 | 5311693 ± 37221 | 0.7 |
| Temperature ($^{\circ}\text{C}$) | 40 | 3.74 ± 0.0 | 0.0 | 4847382 ± 22402 | 0.5 |
| | 30 | 3.66 ± 0.01 | 0.3 | 5942434 ± 37105 | 0.6 |
| Wavelength ( nm) | 242 | 3.7 ± 0.0 | 0.0 | 4847382 ± 22402 | 0.5 |
| | 244 | 3.76 ± 0.02 | 0.5 | 5311693 ± 37221 | 0.7 |
| (c) System suitability |  |  |  |  |  |
| Parameter | Acceptance Criteria | | | Mean ± SD | RSD (%) |
| Retention Time (RT) | Consistent (within $\pm 2\%$) | | | 3.75 ± 0.01 | 0.3 |
| Peak area | %RSD ≤ 2.0 | | | 7775932 ± 15961 | 0.2 |
| Number of Theoretical Plates (N) | > 2000 (depends on column) | | | 3087 ± 13 | 0.4 |
| Tailing Factor (T) | ≤ 2.0 | | | 1.2 ± 0.0 | 0.0 |
| Resolution (Rs) | ≥ 2.0 (between critical peaks) | | | 8.3 ± 0.6 | 0.9 |
| Capacity Factor (k') | 2<k'<10 | | | 3.6 ± 0.0 | 0.3 |

#### 3.5.4. Precision

The method demonstrated excellent precision, confirmed through intra-day and inter-day analyses. Intra-day repeatability, assessed from six replicate injections of BUd (200 µg/mL) within a single day, yielded a concentration of 207 ± 3=µg/mL with a % RSD of 1.3 %. Inter-day precision, evaluated over two consecutive days under the same conditions, resulted in 210 ± 3 µg/mL and a % RSD of 1.4 %. Both values fall well below the 2 % threshold recommended by ICH guidelines, confirming the method’s reproducibility for routine use and long-term studies [34,35].

#### 3.5.5. Sensitivity

The sensitivity of the method was established through both signal-to-noise ratio and statistical calculations, yielding a LOD of 0.04 µg/mL and an LOQ of 1.2 µg/mL. These threshold levels are also stated in the calibration curve **(Fig. 6(a)).** These values indicate that the method can detect trace amounts of BUd, a critical requirement for early-stage drug release studies and degradation profiling, where drug concentrations may fall below standard detection thresholds. The ability to reliably detect and quantify low concentrations also enhances the method’s utility for pharmacokinetic sampling, drug release kinetics, and stability-indicating assays [14,35].

#### 3.5.6. Robustness

Robustness was established by introducing minor, deliberate variations in flow rate, injection volume, column temperature, and detection wavelength **(Table 3(b))**. These changes caused negligible shifts in retention time (% RSD < 0.6%) and peak area (% RSD < 2.0%), confirming method reliability under typical operational fluctuations. For instance, increasing the flow rate from 0.34 to 0.35 mL/min resulted in a marginal shift in retention time from 3.75 to 3.65 ± 0.01=min, with corresponding peak area variation within acceptable limits. No significant changes were observed in peak symmetry or resolution. These results highlight the method’s robustness and operational tolerance, making it suitable for routine use under standard laboratory conditions without risk of analytical drift [15,34].

#### 3.5.7. System Suitability

System suitability testing showed high repeatability at 3.75 ± 0.01=min, with a peak area % RSD of 0.2 %, confirming analytical repeatability **(Table 3(c))**. The theoretical plate count (3088 ± 13) exceeded the recommended minimum of 2000, while the tailing factor was 1.22 ± 0.00 (≥ 2.0), indicating efficient separation and peak symmetry. Moreover, the resolution between critical peaks was 8.3=±=0.7 (≥2.0), and the capacity factor (k’) was 3.62 ± 0.01, both falling within the optimal range for reversed-phase separations. These parameters confirm the system’s operational readiness and the method’s reproducibility under optimized conditions [38].

### 3.6. Application of the validated RP-HPLC method to nanoparticle formulation

#### 3.6.1. Forced degradation studies

Under acidic conditions **(Fig. 7(b) and (c))**, the BUd NPs retained only 48.7 ± 19.3 µg/mL from an initial 209.3 ± 10.5 µg/mL, which translates to roughly 77 % degradation. In comparison, the free drug undergo a moderate degradation labile because of hydrolysis and maintained 102.3 ± 8.6 µg/mL. The substantial degradation observed in both formulations was expected, given the documented sensitivity of PLGA esters to acidic environments and the drug’s proneness to hydrolytic degradation [46,47]. Although the Bud NPs degraded more than the free drug, they still preserved a measurable amount of drug content, implying that the PLL-HA layered structure provided some degree of buffering and steric protection. The statistically significant difference between treated and untreated NPs (p < 0.05) further supports the analytical method’s capacity to indicate stability.

Exposure to alkaline conditions **(Fig. 7(b) and (d))** resulted in the most pronounced degradation among all stress tests. Bud-NPs retained only 22.8 ± 15.2 µg/mL from their initial 118.7 ± 2.8 µg/mL, translating to roughly 80.8 % degradation. However, by comparison, the free drug underwent severe degradation to retain 59.7 ± 2.6 µg/mL drug, indicates considerable instability. The pronounced degradation of BUd NPs under alkaline conditions confirms their base labile nature, primarily because of PLGA hydrolysis that compromises matrix integrity and accelerates drug release. While some drug persisted, likely protected by the PLL and HA layers, the overall loss underscores the critical need for strict pH control during storage and handling [37,47].

Thermal stress **(Fig. 7(b) and (e))** had minimal impact on both the formulation and the drug. BUd NPs retained 117.9 ± 37.9 µg/mL compared to 118.7 ± 2.7 µg/mL in controls, and free BUd remained stable at 203.2 ± 7.8 µg/mL versus 201.4 ± 14.8 µg/mL. Both BUd NPs and free BUd exhibited considerable thermal stability under hydrated conditions, with negligible drug loss and no apparent compromise to the PLGA matrix. These findings suggest the formulation is robust and suitable for storage and transport under moderate temperature conditions [34].

Oxidative degradation **(Fig. 7(b) and (f))** significantly affected BUd-NPs, with only 32.3 ± 20.1 µg/mL remaining from 119.9 ± 23.4 µg/mL (73 % loss), whereas the pure drug retained 190.2 ± 17.1 µg/mL. This differential response reflects the susceptibility of PLGA and surface polymers to peroxidation and oxidative cleavage. These findings are relevant for IBD, where reactive oxygen species are abundant, underscoring the need for oxidative stabilizers in future formulations [34].

Photolytic degradation **(Fig. 7(b) and (g))** resulted in severe loss of BUd from both systems. BUd NPs retained 20.0 ± 9.3 µg/mL, while free BUd dropped to 34.3 ± 0.2 µg/mL, each reflecting 81 % degradation. The hydrated nanoparticle state likely increased permeability, accelerating drug exposure to UV-induced damage. Similarly, photostatic degradation **(Fig. 7(b) and (h))** under daylight exposure reduced BUd NP content to 29.4 ± 5.0 µg/mL, in contrast to 112.0 ± 11.1 µg/mL in controls (74 % loss). These results confirm the loss of photo protective functionality upon hydration and highlight the critical importance of post-reconstitution handling guidelines and light-protective packaging [48].

#### 3.6.2. Determination of drug loading in nanoparticles

Quantification of BUd in BUd NPs yielded a drug content of 169.9 µg/mg of NPs, closely aligning with the theoretical loading of 200 µg/mg. The chromatographic analysis exhibited a sharp, well-resolved peak with no interfering signals, confirming the method’s selectivity for accurately detecting the encapsulated drug within the nanoparticle matrix. The minimal deviation between measured and expected values indicates consistent drug incorporation and negligible loss during formulation. These findings not only confirm the successful loading of BUd into the NPs, but also demonstrate that the drug remained chemically stable throughout the fabrication and lyophilization process, an essential requirement for ensuring therapeutic integrity during storage and subsequent application.

#### 3.6.3. In vitro drug release studies from BUd NPs

The in vitro dissolution/release behaviour of free BUd and BUd encapsulated in NPs (BUd NPs) was evaluated under simulated gastrointestinal conditions relevant to the IBD environment. Data **(Fig. 8(b))** indicate that free BUd undergoes an extremely rapid and nearly complete release, with over 94 % diffusing out in less than 4 h, regardless of pH. This rapid dissolution and diffusion through the dialysis membrane likely accounts for the swift systemic absorption, which could limit the drug’s therapeutic effectiveness in the colon. In contrast, the BUd NPs demonstrated a markedly controlled and sustained release profile throughout the pH transitions, mimicking gastric, intestinal, and colonic environments. Only about 7.6 % of the encapsulated drug was released in the acidic gastric phase (pH 1.8), suggesting substantial nanoparticle stability under these conditions. As the medium’s pH increased, cumulative release from the NPs rose gradually, reaching approximately 15.6 % at 24 h and 20.6 % at 48 h. This sustained release aligns with the structural properties of the PLGA matrix and the surface coatings of PLL and HA, which collectively regulate swelling, degradation, and drug diffusion in a pH-dependent manner. Such a time-dependent release mechanism, involving both diffusion and degradation, is characteristic of PLGA-based NPs and is helpful for localized IBD therapy. By prolonging BUd exposure specifically in the colon, the formulation enhances anti-inflammatory efficacy while minimizing systemic corticosteroid side effects. Overall, these findings support the potential of BUd NPs as an effective oral delivery system for targeted treatment of colonic inflammation in IBD patients [4,42,49].

### 3.7. Greenness profile assessment of the proposed method

The greenness of the developed RP-HPLC method was evaluated using NEMI, GAPI, AGREE, and BAGI **(Fig. 9(a–d))**. The NEMI pictogram **(Fig. 9a)** showed three green quadrants (non-PBT solvent, non-corrosive pH, and low waste generation <10 mL per run), while the hazardous waste quadrant remained white due to the use of acetonitrile. This outcome indicates broad compliance with basic GAC principles, though solvent choice still restricts complete greenness [32,50]. The GAPI pictogram **(Fig. 9b)** presented a balanced but cautious profile. Red zones were observed for sample collection (criterion 1) and solvent use (criterion 7), reflecting off-line sampling and reliance on acetonitrile. Three green fields— preservation (2), low solvent amount (<10 mL, criterion 9), and minimal occupational hazard (13)—highlighted favourable features. The remaining parameters appeared yellow, indicating moderate impact across sample preparation, energy consumption, and waste treatment. Overall, this assessment suggests the method is environmentally acceptable, with solvent replacement and workflow integration offering the greatest opportunities for improvement [51]. The AGREE analysis **(Fig. 9c)** yielded a score of **0.69**, with strengths stemming from short run time (<5 min), buffer-free composition, and reduced solvent volumes. Penalties were assigned for solvent selection and the off-line analytical format, but no severe violations were observed. The clock visualization confirmed that most principles fell within the green– yellow range [51]. The BAGI index **(Fig. 9d)** returned a score of 70/100, with high points awarded for use of common reagents, simple preparation, absence of preconcentration, and semi-automation. Lower scores were linked to single-analyte determination and moderate sample volumes, highlighting potential improvements through multiplexing or miniaturization [51,52]. Together, the four tools consistently indicate that the developed RP-HPLC method offers a greener alternative to conventional chromatographic protocols. Short analysis time, low waste generation, and simple preparation underpin its sustainability advantages, while the continued reliance on acetonitrile remains the most critical limitation to achieving a fully green profile.

**Figure 9.**
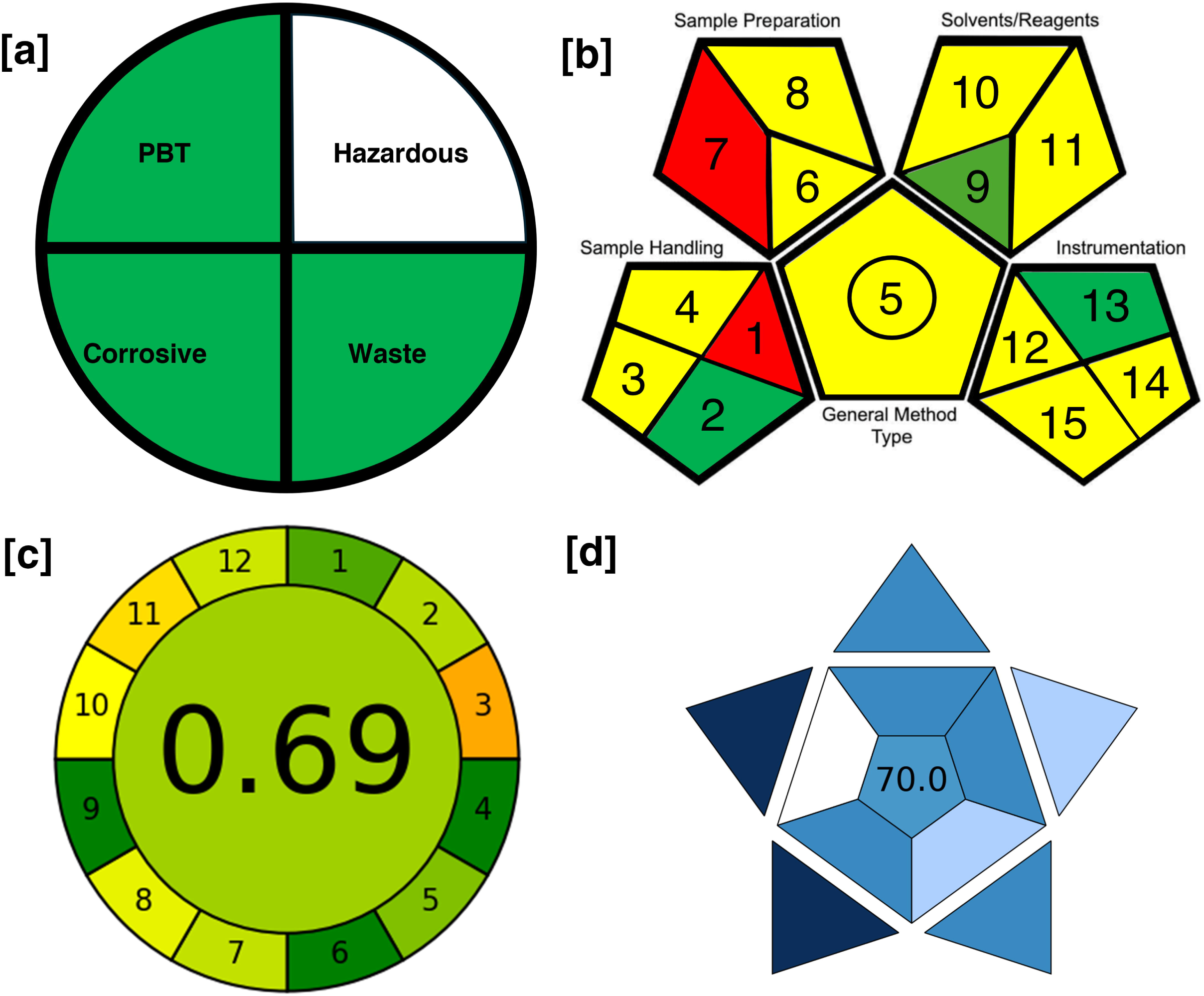
Greenness assessment of the developed RP-HPLC method using (a) NEMI, (b) GAPI, (c) AGREE, and (d) BAGI.

## 4. Conclusion

This study reports the first validated RP-HPLC–UV method specifically developed for quantifying budesonide encapsulated within poly(lactic-co-glycolic acid)-based, layer-by-layer (LbL) coated nanoparticles intended for colonic drug delivery. Employing a response surface methodology–central composite design approach, the method was optimized to use a rapid, isocratic, buffer-free acetonitrile/water (80:20, v/v) mobile phase, enabling complete separation within 5 min at a flow rate of 0.34 mL/min and a 10 µL injection volume. Validation confirmed compliance with ICH Q2(R2) criteria, including linearity, accuracy, precision, sensitivity, specificity, robustness, and system suitability, confirming its suitability for both formulation development and quality control.

In addition to being analytically robust, the method demonstrated a favorable greenness profile, with minimal solvent consumption, reduced analysis time, and the absence of toxic buffers. This was further supported by GAPI and NEMI assessments, which verified the method’s low environmental impact while maintaining high analytical performance. Application to forced degradation studies showed that the LbL coating markedly improved budesonide stability under thermal and photostatic stress while providing measurable protection against hydrolytic, oxidative, and photolytic degradation. In vitro release profiling under simulated gastrointestinal pH conditions demonstrated a sustained, pH-responsive release (20.6% over 48 h), consistent with targeted colonic delivery objectives. Overall, this analytical platform offers a green, stability-indicating, and versatile solution for comprehensive performance assessment of advanced nanoparticulate drug delivery systems. By balancing environmental sustainability with functionality, it not only supports formulation development and quality control but also advances the broader goal of implementing greener regulatory methods in pharmaceutical analysis and clinical translation.

## Acknowledgements

This work was supported by a grant from the Science Foundation Ireland (SFI) and the European Regional Development Fund (ERDF) under Grant number 13/RC/2073_P2. The authors acknowledge the facilities of the Anatomy Imaging and Microscopy Facility at the University of Galway and the technical assistance of Dr Éadaoin Timmins https://imaging.universityofgalway.ie/imaging/).

## CRediT authorship contribution statement

**Giriprasath Ramanathan:** Conceptualisation, Methodology, Investigation, Validation, Data curation, Software, Formal analysis, Writing – Original Draft, Writing – review & editing, Visualisation. **Masroora Hassan:** Conceptualisation, Investigation, Formal analysis, Writing – Review & Editing. **Yury Rochev:** Conceptualisation, Writing – review & editing, Resources, Supervision, Validation, Funding acquisition, Project administration.

