## Supplementary Material for "Greenness-Assessed RP-HPLC Method for Budesonide Quantification in Layer-by-Layer Polymeric Nanoparticles: Validation, Stability, and Controlled Release in Colonic Delivery" (from another version of this preprint: version 1, 2025-08-15)

**Optimisation and validation of a rapid RP-HPLC method for budesonide quantification from polymeric nanoparticles with application to stability and drug release studies**

Giriprasath Ramanathan* ^a b^, Masroora Hassan ^a b^, Yury Rochev* ^a b^

^a^ School of Physics, University of Galway, Galway, Ireland

^b^ CÚRAM, Research Ireland Centre for Medical Devices, University of Galway, Galway *H91 W2TY,* Ireland

*****Corresponding authors

Dr. Yury Rochev,

Dr. Giriprasath Ramanathan,

**Table S1. Experimental design and response data from Phase II of the RSM-CCD study. Variables included mobile phase composition (A), flow rate (B), and solvent used for drug dissolution (C). Responses recorded were retention time (R_1_), peak area (R_2_), and calculated budesonide concentration (R_2a_), based on the optimized calibration curve. Values are reported as mean ± SD (n = 3).**

|  | **DOE Exp. No 1.** | | | | | |
| --- | --- | --- | --- | --- | --- | --- |
| **Run order** | **Numerical variables** | | **Categorical Variables** | **Response (R)** | | |
|  | **A:MP (MeOH:H_2_O)** | **B: FR (mL/min)** | **C: DD**  **(1 mL)** | **R_1_: Rt (min)** | **R_2_: PA (AU)** | **R_2a_: Concentration (µg/mL)** |
| 1 | 70:30 | 0.375 | MeOH | 6.68 ± 0.01 | 5040769 ± 21915 | 89.5 ± 0.4 |
| 2 | 70:30 | 0.375 | EtOH | 6.56 ± 0.01 | 4754633 ± 3030 | 84.4 ± 0.1 |
| 3 | 80:20 | 0.375 | MeOH | 4.28 ± 0.00 | 5071258 ± 10221 | 90.1 ± 0.2 |
| 4 | 75:25 | 0.4 | ACN | 4.79 ± 0.00 | 5141230 ± 3046 | 91.3 ± 0.1 |
| 5 | 70:30 | 0.425 | EtOH | 5.79 ± 0.01 | 4188915 ± 10303 | 74.2 ± 0.2 |
| 6 | 80:20 | 0.425 | EtOH | 3.73 ± 0.00 | 4332455 ± 41856 | 76.9 ± 0.6 |
| 7 | 75:25 | 0.4 | EtOH | 4.72 ± 0.00 | 4522290 ± 8979 | 80.3 ± 0.2 |
| 8 | 75:25 | 0.4 | EtOH | 4.71 ± 0.00 | 4523820 ± 8979 | 80.3 ± 0.2 |
| 9 | 75:25 | 0.4 | MeOH | 4.81 ± 0.00 | 4728500 ± 1409 | 84.0 ± 0.0 |
| 10 | 75:25 | 0.4 | ACN | 4.79 ± 0.00 | 5137410 ± 3046 | 91.3 ± 0.1 |
| 11 | 75:25 | 0.4 | MeOH | 4.81 ± 0.00 | 4732210 ± 1409 | 84.0 ± 0.0 |
| 12 | 70:30 | 0.425 | MeOH | 5.89 ± 0.00 | 4439957 ± 12316 | 78.8 ± 0.2 |
| 13 | 80:20 | 0.425 | MeOH | 3.78 ± 0.00 | 4484710 ± 3776 | 79.6 ± 0.1 |
| 14 | 70:30 | 0.425 | ACN | 5.98 ± 0.00 | 4823560 ± 9928 | 85.7 ± 0.2 |
| 15 | 70:30 | 0.375 | ACN | 6.85 ± 0.01 | 5474040 ± 25243 | 97.3 ± 0.5 |
| 16 | 80:20 | 0.425 | ACN | 3.77 ± 0.00 | 4875680 ± 17526 | 86.6 ± 0.3 |
| 17 | 75:25 | 0.4 | ACN | 4.79 ± 0.00 | 5134920 ± 3046 | 91.2 ± 0.1 |
| 18 | 75:25 | 0.4 | MeOH | 4.81 ± 0.00 | 4729890 ± 1409 | 84.0 ± 0.0 |
| 19 | 75:25 | 0.4 | EtOH | 4.71 ± 0.00 | 4523170 ± 8979 | 80.3 ± 0.7 |
| 20 | 80:20 | 0.375 | EtOH | 4.22 ± 0.00 | 4878130 ± 2238 | 86.7 ± 0.0 |
| 21 | 80:20 | 0.375 | ACN | 4.28 ± 0.00 | 5529250 ± 16873 | 98.3 ± 0.3 |
| 22 | 75:25 | 0.4 | EtOH | 4.72 ± 0.00 | 4523860 ± 8979 | 80.3 ± 0.2 |
| 23 | 75:25 | 0.375 | EtOH | 5.02 ± 0.01 | 4876940 ± 54982 | 86.7 ± 0.0 |
| 24 | 80:20 | 0.4 | EtOH | 3.96 ± 0.00 | 4532580 ± 30294 | 80.4 ± 0.5 |
| 25 | 75:25 | 0.4 | EtOH | 4.72 ± 0.00 | 4545570 ± 8979 | 80.7 ± 0.2 |
| 26 | 75:25 | 0.425 | EtOH | 4.44 ± 0.00 | 4220460 ± 32556 | 74.8 ± 0.6 |
| 27 | 75:25 | 0.4 | EtOH | 4.72 ± 0.00 | 4525520 ± 8979 | 80.3 ± 0.7 |
| 28 | 75:25 | 0.375 | ACN | 5.11 ± 0.01 | 5478190 ± 1325 | 97.4 ± 0.0 |
| 29 | 75:25 | 0.4 | MeOH | 4.81 ± 0.00 | 4729460 ± 1409 | 84.0 ± 0.0 |
| 30 | 80:20 | 0.4 | MeOH | 4.02 ± 0.00 | 4773690 ± 9674 | 84.8 ± 0.2 |
| 31 | 75:25 | 0.4 | ACN | 4.79 ± 0.00 | 5143250 ± 3046 | 91.4 ± 0.1 |
| 32 | 70:30 | 0.4 | MeOH | 6.26 ± 0.00 | 4723901 ± 14658 | 83.9 ± 0.2 |
| 33 | 75:25 | 0.4 | ACN | 4.79 ± 0.00 | 5141170 ± 3046 | 91.3 ± 0.1 |
| 34 | 70:30 | 0.4 | EtOH | 6.14 ± 0.00 | 4475556 ± 51254 | 79.4 ± 0.1 |
| 35 | 70:30 | 0.4 | ACN | 6.39 ± 0.00 | 5103647 ± 9637 | 90.5 ± 0.2 |
| 36 | 75:25 | 0.425 | MeOH | 4.52 ± 0.00 | 4452956 ± 5258 | 79.0 ± 0.1 |
| 37 | 75:25 | 0.425 | ACN | 4.51 ± 0.00 | 4836040 ± 6192 | 85.9 ± 0.1 |
| 38 | 80:20 | 0.4 | ACN | 4.01 ± 0.00 | 5191418 ± 8802 | 92.2 ± 0.1 |
| 39 | 75:25 | 0.375 | MeOH | 5.13 ± 0.01 | 5042065 ± 2535 | 89.6 ± 0.1 |
| 40 | 75:25 | 0.4 | MeOH | 4.81 ± 0.00 | 4731130 ± 1409 | 84.0 ± 0.0 |
| 41 | 75:25 | 0.4 | MeOH | 4.80 ± 0.00 | 4728900 ± 1409 | 84.0 ± 0.0 |
| 42 | 75:25 | 0.4 | ACN | 4.79 ± 0.00 | 5140830 ± 3046 | 91.3 ± 0.1 |

**Table S2. Experimental design and response data from Phase II of the RSM-CCD study. Variables included flow rate (D), injection volume (E), and drug dissolution solvent (F). Responses recorded were retention time (R₃), peak area (R₄), and calculated budesonide concentration (R_4a_), based on the optimized calibration curve. Values are reported as mean ± SD (n = 3).**

| **DOE Exp. No 2.** | | | | | | |
| --- | --- | --- | --- | --- | --- | --- |
| **Run order** | **Numerical variables** | | **Categorical Variables** | **Response (R)** | | |
|  | **D: FR (mL/min)** | **E: IV (µL)** | **F: DD**  **(1 mL)** | **R_3_: Rt (min)** | **R_4_: PA (AU)** | **R_4a_: Concentration (µg/mL)** |
| 1 | 0.36 | 5 | ACN | 3.54 ± 0.00 | 1565140 ± 193020.28 | 29.2 ± 3.6 |
| 2 | 0.36 | 15 | MeOH | 3.59 ± 0.00 | 3719690 ± 126271.27 | 70.0 ± 2.4 |
| 3 | 0.34 | 15 | EtOH | 3.77 ± 0.01 | 5132656 ± 179623.87 | 96.7 ± 3.4 |
| 4 | 0.36 | 5 | EtOH | 3.54 ± 0.01 | 1310628 ± 58742.86 | 24.4 ± 1.1 |
| 5 | 0.35 | 10 | ACN | 3.65 ± 0.01 | 3539444 ± 1071076.45 | 66.6 ± 20.3 |
| 6 | 0.35 | 10 | MeOH | 3.66 ± 0.01 | 3649879 ± 621619.11 | 68.7 ± 11.8 |
| 7 | 0.35 | 10 | EtOH | 3.66 ± 0.02 | 3137176 ± 1190856.37 | 58.9 ± 22.5 |
| 8 | 0.35 | 10 | ACN | 3.65 ± 0.01 | 3113583 ± 1071076.45 | 58.5 ± 20.3 |
| 9 | 0.35 | 10 | MeOH | 3.66 ± 0.01 | 3639191 ± 621619.11 | 68.4 ± 11.8 |
| 10 | 0.34 | 15 | ACN | 3.76 ± 0.02 | 5311693 ± 537221.88 | 100.1 ± 10.2 |
| 11 | 0.35 | 10 | ACN | 3.66 ± 0.01 | 3346724 ± 1071076.45 | 62.9 ± 20.3 |
| 12 | 0.34 | 15 | MeOH | 3.78 ± 0.00 | 5044888 ± 123993.28 | 95.0 ± 2.3 |
| 13 | 0.36 | 5 | MeOH | 3.54 ± 0.01 | 1251315 ± 65977.86 | 23.2 ± 1.3 |
| 14 | 0.35 | 10 | EtOH | 3.66 ± 0.02 | 3194821 ± 1190856.37 | 60.0 ± 22.5 |
| 15 | 0.34 | 5 | ACN | 3.74 ± 0.00 | 2670394 ± 81834.00 | 50.1 ± 1.5 |
| 16 | 0.34 | 5 | EtOH | 3.75 ± 0.00 | 2566682 ± 115571.33 | 48.1 ± 2.2 |
| 17 | 0.36 | 15 | EtOH | 3.77 ± 0.01 | 5132656 ± 179623.87 | 96.7 ± 3.4 |
| 18 | 0.34 | 5 | MeOH | 3.75 ± 0.01 | 2073848 ± 173441.31 | 38.8 ± 3.2 |
| 19 | 0.36 | 15 | ACN | 3.57 ± 0.00 | 4528289 ± 131939.07 | 85.3 ± 2.5 |
| 20 | 0.35 | 10 | EtOH | 3.61 ± 0.02 | 2922581 ± 1190856.37 | 54.9 ± 22.5 |
| 21 | 0.35 | 10 | MeOH | 3.66 ± 0.01 | 3416194 ± 621619.11 | 64.3 ± 11.7 |
| 22 | 0.35 | 10 | MeOH | 3.66 ± 0.01 | 4734565 ± 621619.11 | 89.2 ± 11.7 |
| 23 | 0.34 | 10 | EtOH | 3.76 ± 0.02 | 4801677 ± 447860.57 | 90.5 ± 8.4 |
| 24 | 0.34 | 10 | ACN | 3.74 ± 0.00 | 4847382 ± 522402.51 | 91.3 ± 9.8 |
| 25 | 0.35 | 5 | EtOH | 3.64 ± 0.01 | 1240587 ± 46224.53 | 23.1 ± 0.8 |
| 26 | 0.35 | 10 | ACN | 3.64 ± 0.01 | 5271123 ± 1071076.45 | 99.4 ± 20.3 |
| 27 | 0.35 | 15 | EtOH | 3.66 ± 0.02 | 5327457 ± 129936.00 | 100.4 ± 2.5 |
| 28 | 0.35 | 10 | ACN | 3.64 ± 0.01 | 5290105 ± 1071076.45 | 99.7 ± 20.3 |
| 29 | 0.35 | 5 | ACN | 3.64 ± 0.00 | 1260778 ± 235738.30 | 23.5 ± 4.5 |
| 30 | 0.35 | 10 | ACN | 3.64 ± 0.01 | 5258087 ± 1071076.45 | 99.1 ± 20.3 |
| 31 | 0.36 | 10 | ACN | 3.55 ± 0.01 | 2620036 ± 950330.71 | 49.2 ± 17.9 |
| 32 | 0.36 | 10 | EtOH | 3.56 ± 0.00 | 3008092 ± 80690.04 | 56.5 ± 1.5 |
| 33 | 0.35 | 5 | MeOH | 3.65 ± 0.00 | 1956529 ± 81434.15 | 36.6 ± 1.5 |
| 34 | 0.36 | 10 | MeOH | 3.57 ± 0.00 | 2698530 ± 128599.59 | 50.7 ± 2.4 |
| 35 | 0.35 | 15 | MeOH | 3.68 ± 0.01 | 5115437 ± 488526.00 | 96.4 ± 9.2 |
| 36 | 0.35 | 10 | MeOH | 3.66 ± 0.01 | 4797382 ± 621619.11 | 90.4 ± 11.8 |
| 37 | 0.35 | 15 | ACN | 3.66 ± 0.01 | 5942434 ± 537105.68 | 112.1 ± 10.2 |
| 38 | 0.35 | 10 | EtOH | 3.67 ± 0.02 | 5305058 ± 1190856.37 | 100.0 ± 22.5 |
| 39 | 0.35 | 10 | MeOH | 3.65 ± 0.01 | 4504788 ± 621619.11 | 84.9 ± 11.8 |
| 40 | 0.34 | 10 | MeOH | 3.77 ± 0.00 | 3900528 ± 122251.41 | 73.4 ± 2.3 |
| 41 | 0.35 | 10 | EtOH | 3.65 ± 0.02 | 5411512 ± 1190856.37 | 102.0 ± 22.5 |
| 42 | 0.35 | 10 | EtOH | 3.65 ± 0.02 | 4999242 ± 1190856.37 | 94.2 ± 22.5 |

**Table S3. ANOVA summary for the response surface quadratic models evaluating (a) retention time (R₃) and (b) peak area (R₄) in Phase I of the CCD-based DOE experiment. Main effects, interactions, and model fit terms are reported with corresponding F- and p-values.**

| (a) Retention time (Rt) | | | | | |
| --- | --- | --- | --- | --- | --- |
| Source | Sum of Squares | df | Mean Square | F-value | p-value |
| Block | 0.1253 | 1 | 0.1253 |  |  |
| Model | 26.65 | 11 | 2.42 | 3919.94 | < 0.0001 |
| A-A | 23.37 | 1 | 23.37 | 37810.97 | < 0.0001 |
| B-B | 1.81 | 1 | 1.81 | 2931.80 | < 0.0001 |
| C-C | 0.0955 | 2 | 0.0477 | 77.21 | < 0.0001 |
| AB | 0.0747 | 1 | 0.0747 | 120.90 | < 0.0001 |
| AC | 0.0300 | 2 | 0.0150 | 24.26 | < 0.0001 |
| BC | 0.0015 | 2 | 0.0007 | 1.19 | 0.3199 |
| A² | 1.06 | 1 | 1.06 | 1710.39 | < 0.0001 |
| B² | 0.0028 | 1 | 0.0028 | 4.57 | 0.0411 |
| Residual | 0.0179 | 29 | 0.0006 |  |  |
| Lack of Fit | 0.0178 | 17 | 0.0010 | 130.19 | < 0.0001 |
| Pure Error | 0.0001 | 12 | 8.056E-06 |  |  |
| Cor Total | 26.80 | 41 |  |  |  |
| (b) Peak area (PA) | | | | | |
| Source | Sum of Squares | df | Mean Square | F-value | p-value |
| Block | 5.335E+08 | 1 | 5.335E+08 |  |  |
| Model | 4.520E+12 | 11 | 4.109E+11 | 1392.43 | < 0.0001 |
| A-A | 2.305E+10 | 1 | 2.305E+10 | 78.13 | < 0.0001 |
| B-B | 1.675E+12 | 1 | 1.675E+12 | 5675.37 | < 0.0001 |
| C-C | 2.810E+12 | 2 | 1.405E+12 | 4761.60 | < 0.0001 |
| AB | 8.115E+07 | 1 | 8.115E+07 | 0.2750 | 0.6040 |
| AC | 3.397E+09 | 2 | 1.699E+09 | 5.76 | 0.0078 |
| BC | 3.371E+09 | 2 | 1.686E+09 | 5.71 | 0.0081 |
| A² | 1.127E+08 | 1 | 1.127E+08 | 0.3820 | 0.5414 |
| B² | 3.731E+09 | 1 | 3.731E+09 | 12.64 | 0.0013 |
| Residual | 8.558E+09 | 29 | 2.951E+08 |  |  |
| Lack of Fit | 8.231E+09 | 17 | 4.842E+08 | 17.78 | < 0.0001 |
| Pure Error | 3.267E+08 | 12 | 2.723E+07 |  |  |
| Cor Total | 4.529E+12 | 41 |  |  |  |

**Table S4. ANOVA summary for the response surface quadratic models evaluating (a) retention time (R₃) and (b) peak area (R₄) in Phase II of the CCD-based DOE experiment. Main effects, interactions, and model fit terms are reported with corresponding F- and p-values.**

| (a) Retention time (Rt) | | | | | |
| --- | --- | --- | --- | --- | --- |
| Source | Sum of Squares | df | Mean Square | F-value | p-value |
| Block | 0.0016 | 1 | 0.0016 |  |  |
| Model | 0.1669 | 11 | 0.0152 | 21.17 | < 0.0001 |
| A-A | 0.1383 | 1 | 0.1383 | 192.97 | < 0.0001 |
| B-B | 0.0112 | 1 | 0.0112 | 15.55 | 0.0005 |
| C-C | 0.0029 | 2 | 0.0014 | 2.00 | 0.1539 |
| AB | 0.0046 | 1 | 0.0046 | 6.37 | 0.0174 |
| AC | 0.0034 | 2 | 0.0017 | 2.35 | 0.1128 |
| BC | 0.0038 | 2 | 0.0019 | 2.65 | 0.0873 |
| A² | 0.0011 | 1 | 0.0011 | 1.60 | 0.2158 |
| B² | 0.0007 | 1 | 0.0007 | 0.9153 | 0.3466 |
| Residual | 0.0208 | 29 | 0.0007 |  |  |
| Lack of Fit | 0.0189 | 17 | 0.0011 | 7.18 | 0.0006 |
| Pure Error | 0.0019 | 12 | 0.0002 |  |  |
| Cor Total | 0.1893 | 41 |  |  |  |
| (b) Peak area (PA) | | | | | |
| Block | 7.735E+12 | 1 | 7.735E+12 |  |  |
| Model | 5.745E+13 | 11 | 5.223E+12 | 10.62 | < 0.0001 |
| A-A | 6.143E+12 | 1 | 6.143E+12 | 12.49 | 0.0014 |
| B-B | 4.789E+13 | 1 | 4.789E+13 | 97.36 | < 0.0001 |
| C-C | 6.330E+11 | 2 | 3.165E+11 | 0.6435 | 0.5328 |
| AB | 9.634E+10 | 1 | 9.634E+10 | 0.1959 | 0.6614 |
| AC | 1.008E+11 | 2 | 5.040E+10 | 0.1025 | 0.9029 |
| BC | 3.559E+11 | 2 | 1.779E+11 | 0.3617 | 0.6996 |
| A² | 3.357E+11 | 1 | 3.357E+11 | 0.6826 | 0.4154 |
| B² | 1.149E+12 | 1 | 1.149E+12 | 2.34 | 0.1373 |
| Residual | 1.426E+13 | 29 | 4.919E+11 |  |  |
| Lack of Fit | 1.396E+13 | 17 | 8.211E+11 | 32.14 | < 0.0001 |
| Pure Error | 3.065E+11 | 12 | 2.554E+10 |  |  |
| Cor Total | 7.945E+13 | 41 |  |  |  |

**Table S5. Summary of calibration data for BUd (1–200 µg/mL), presented as mean ± SD and % RSD for each concentration point (n = 6).**

| Concentration (µg/mL) | Mean ± SD | %RSD |
| --- | --- | --- |
| 1 | 78070 ± 1336 | 1.7 |
| 5 | 264639 ± 5003 | 1.9 |
| 10 | 555961 ± 8752 | 1.6 |
| 20 | 1032135 ± 10531 | 1.0 |
| 30 | 1536978 ± 21795 | 1.4 |
| 40 | 2118365 ± 26711 | 1.3 |
| 50 | 2718943 ± 38885 | 1.4 |
| 60 | 3208949 ± 55193 | 1.7 |
| 70 | 3808111 ± 46302 | 1.2 |
| 80 | 4214015 ± 72020 | 1.7 |
| 90 | 4814726 ± 68413 | 1.4 |
| 100 | 5842238 ± 101639 | 1.7 |
| 150 | 8323759 ± 148220 | 1.8 |
| 200 | 110000000 ± 187722 | 1.7 |

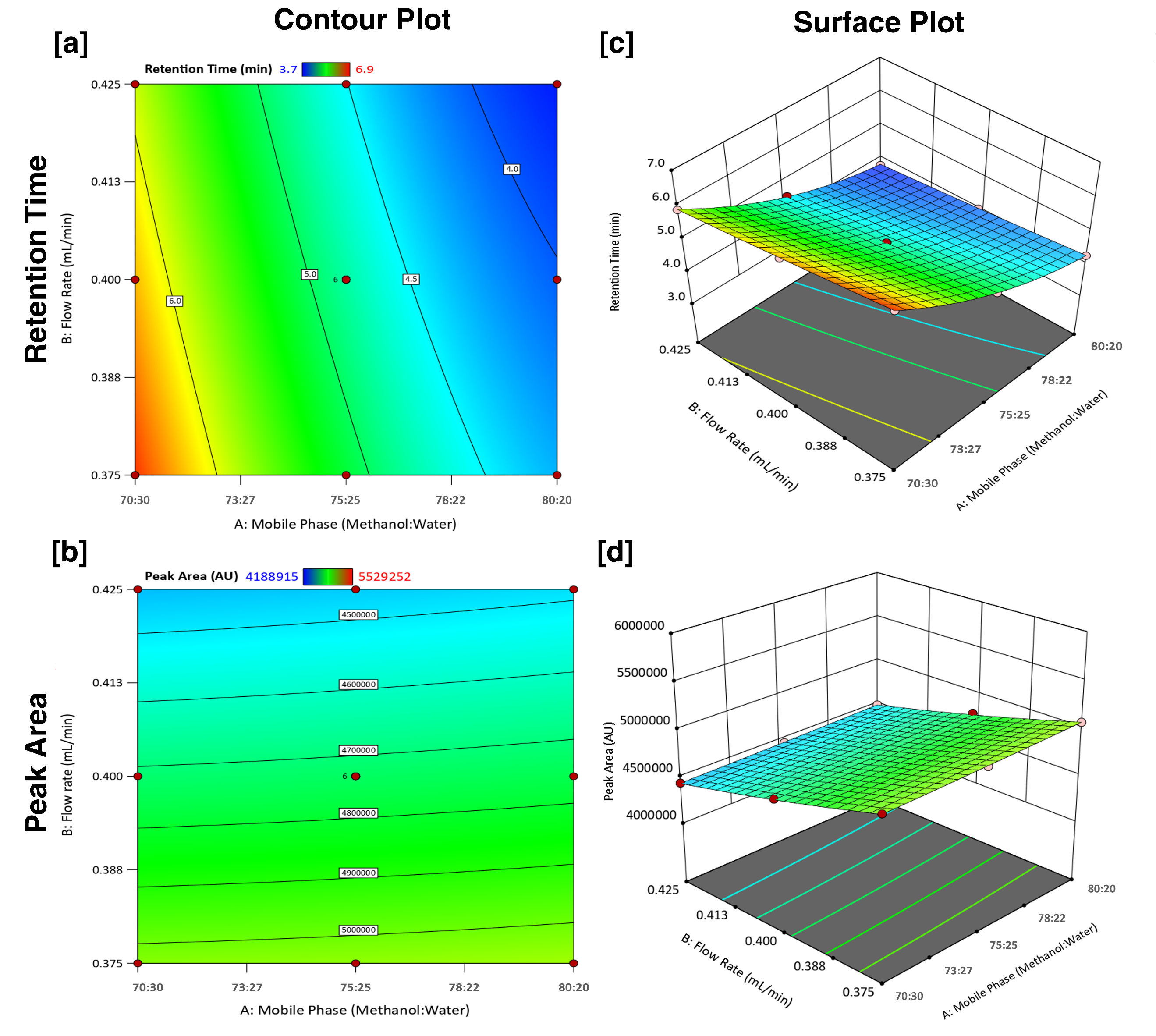

Figure S1. Visualization of variable interactions for Phase I optimization of the RP-HPLC method, with MeOH used as the categorical solvent for drug dissolution. (a, b) Contour plots; and (c, d) 3D surface plots showing the combined effects of mobile phase composition (MeOH:H_2_O at 70:30, 75:25, and 80:20 v/v) and flow rate (0.375, 0.400, and 0.425 mL/min) on retention time and peak area. Experiments were performed in triplicate (n = 3).

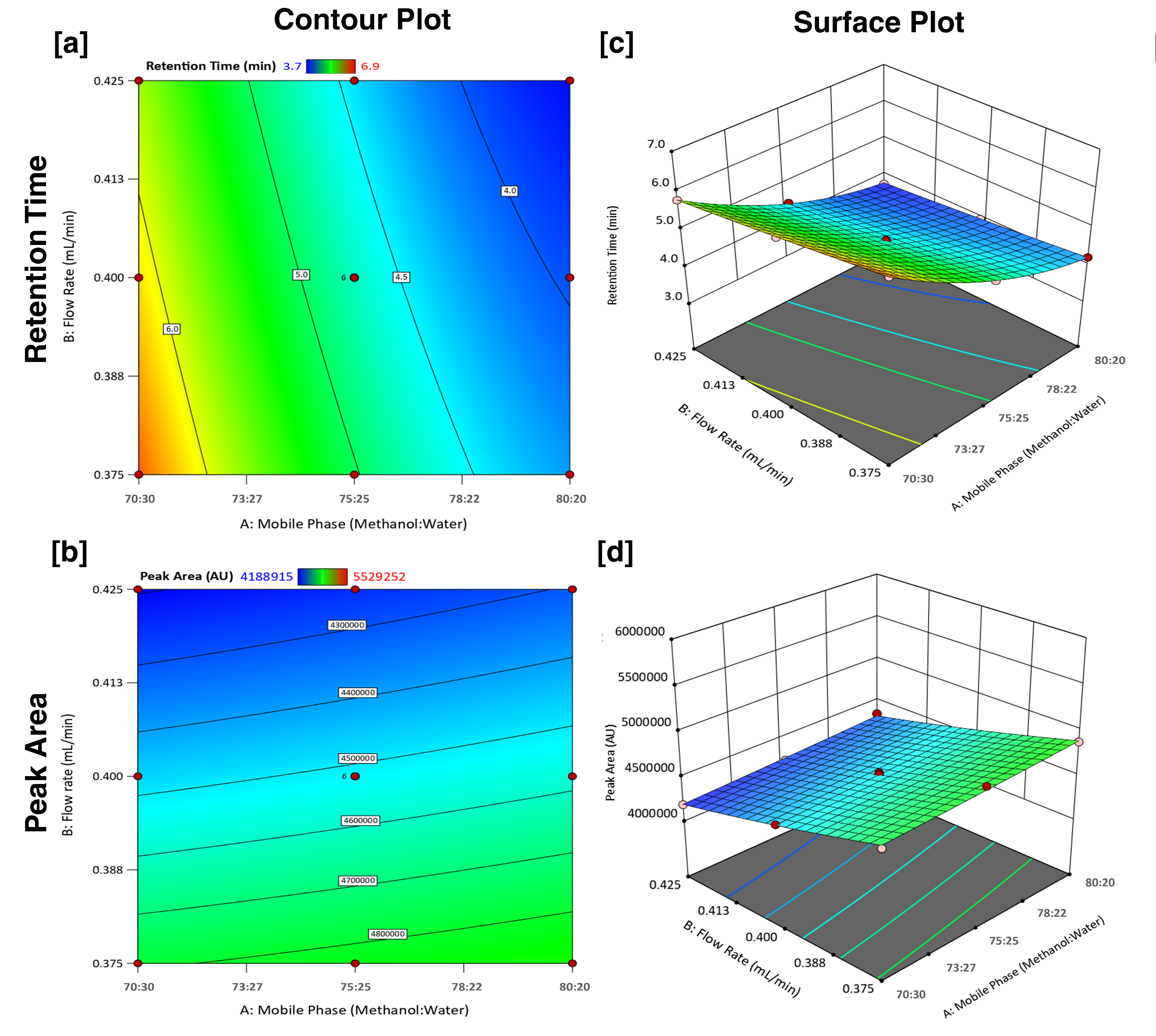

Figure S2. Visualization of variable interactions and model predictions for Phase I optimization of the RP-HPLC method, with EtOH used as the categorical solvent for drug dissolution. (a, b) Contour plots; and (c, d) 3D surface plots showing the combined effects of mobile phase composition (MeOH: H_2_O at 70:30, 75:25, and 80:20 v/v) and flow rate (0.375, 0.400, and 0.425 mL/min) on retention time and peak area. Experiments were performed in triplicate (n = 3).

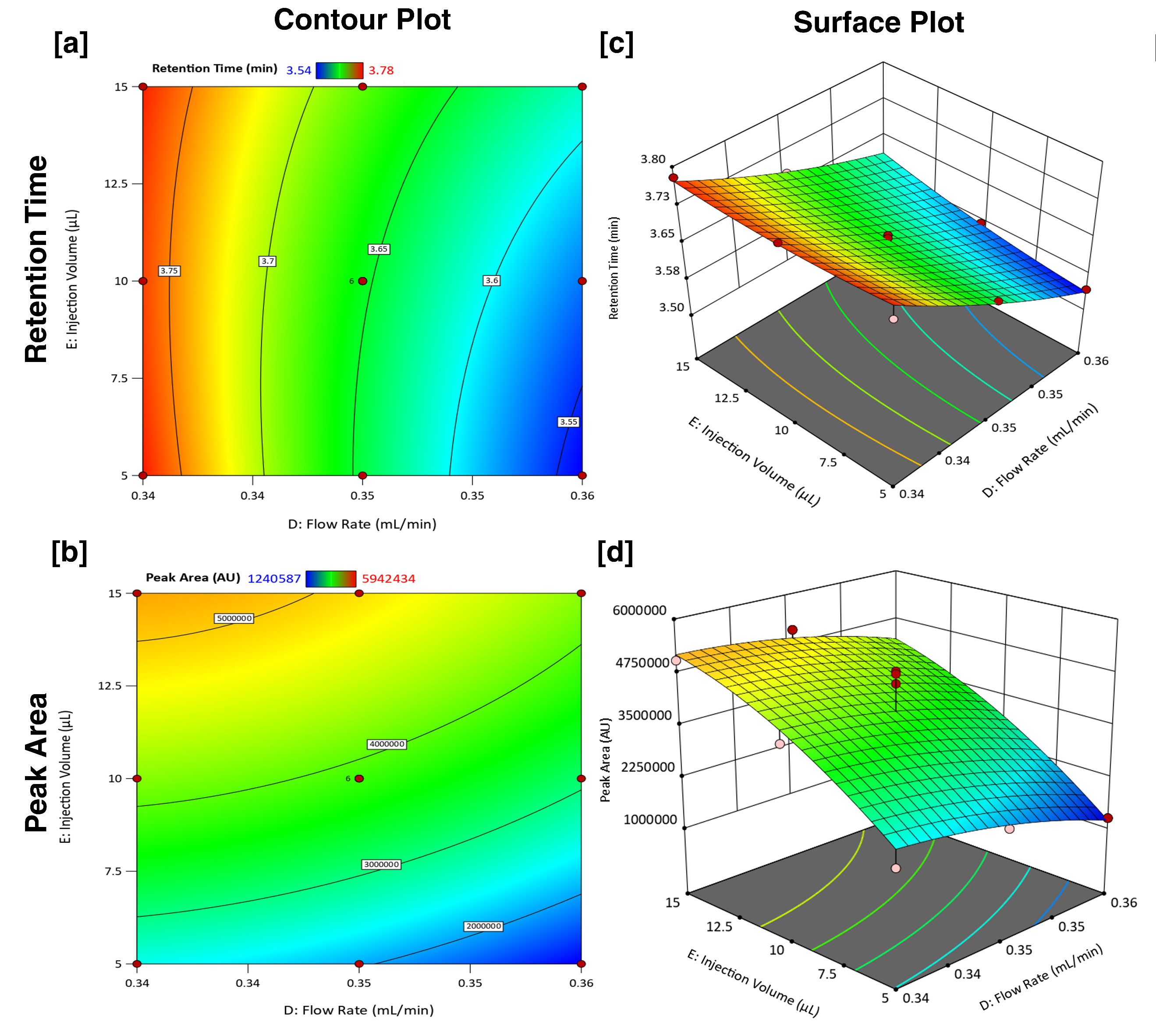

Figure S3. Visualization of variable interactions and model predictions for Phase II optimization of the RP-HPLC method, with MeOH used as the categorical solvent for drug dissolution. (a, b) Contour plots; and (c, d) 3D surface plots; showing the combined effects of flow rate (0.34, 0.35, and 0.36 mL/min) and injection volume (5, 10, and 15 µL) on retention time and peak area. Experiments were performed in triplicate (n = 3).

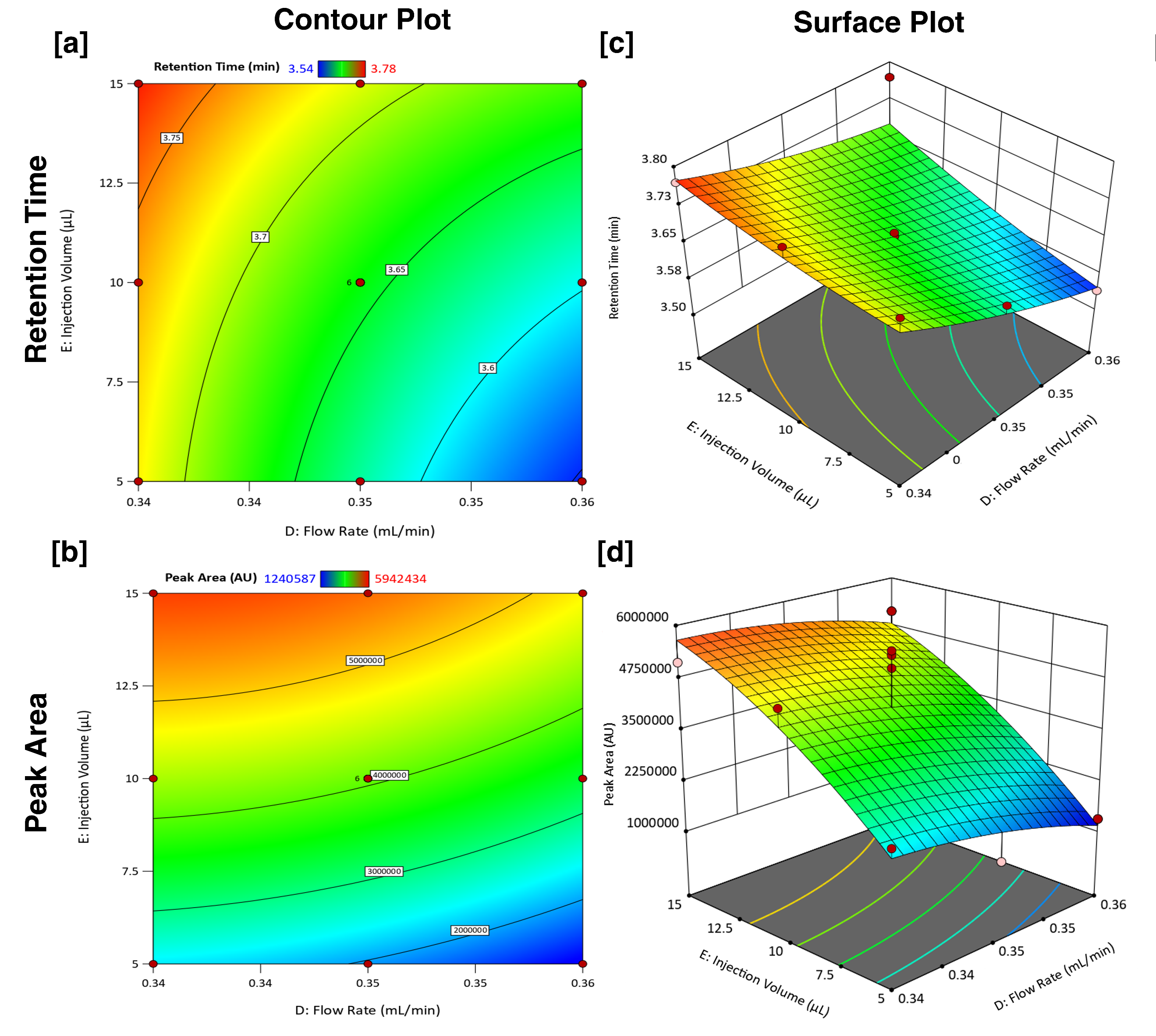

Figure S4. Visualization of variable interactions and model predictions for Phase II optimization of the RP-HPLC method, with EtOH used as the categorical solvent for drug dissolution. (a, b) Contour plots; and (c, d) 3D surface plots; showing the combined effects of flow rate (0.34, 0.35, and 0.36 mL/min) and injection volume (5, 10, and 15 µL) on retention time and peak area. Experiments were performed in triplicate (n = 3).
